# FoxO transcription factors couple the urea cycle and gluconeogenesis by controlling Ass1

**DOI:** 10.64898/2025.12.22.695746

**Authors:** Samia Karkoutly, Yoshinori Takeuchi, Zahra Mehrazad Saber, Duhan Tao, Tsolmon Mendsaikhan, Rika Saikawa, Yuichi Aita, Yuki Murayama, Akito Shikama, Yukari Masuda, Naoya Yahagi

## Abstract

Amino acid catabolism during fasting requires coordinated nitrogen disposal and glucose production, but the transcriptional logic linking the urea cycle to gluconeogenesis remains unclear. Forkhead box O (FoxO) transcription factors are key regulators of fasting metabolism, yet their role in controlling urea cycle genes has not been fully defined. Here we identify FoxOs as direct regulators of hepatic argininosuccinate synthase 1 (Ass1). Because FoxOs often act through Kruppel-like factor 15 (Klf15) in amino acid metabolism, we tested whether Ass1 regulation requires Klf15. Acute hepatic FoxO1/3a knockdown in fasted mice selectively reduced Ass1 expression, lowered blood glucose, and shifted urea-cycle amino acids, with arginine decreased and ornithine increased, even in Klf15-deficient livers. Silencing Ass1 phenocopied these metabolic effects, indicating that Ass1 mediates a key FoxO-dependent branch of fasting adaptation. Mechanistically, we mapped a functional FoxO-binding element within an upstream *Ass1* enhancer: FoxO1/3a activated the enhancer in reporter assays, EMSA confirmed binding, and in vivo luciferase imaging and liver ChIP demonstrated fasting-inducible enhancer activity and FoxO occupancy. Collectively, these findings establish a FoxO-Ass1 axis that couples ureagenesis to gluconeogenesis and supports metabolic flexibility during fasting.

**Graphical abstract:** 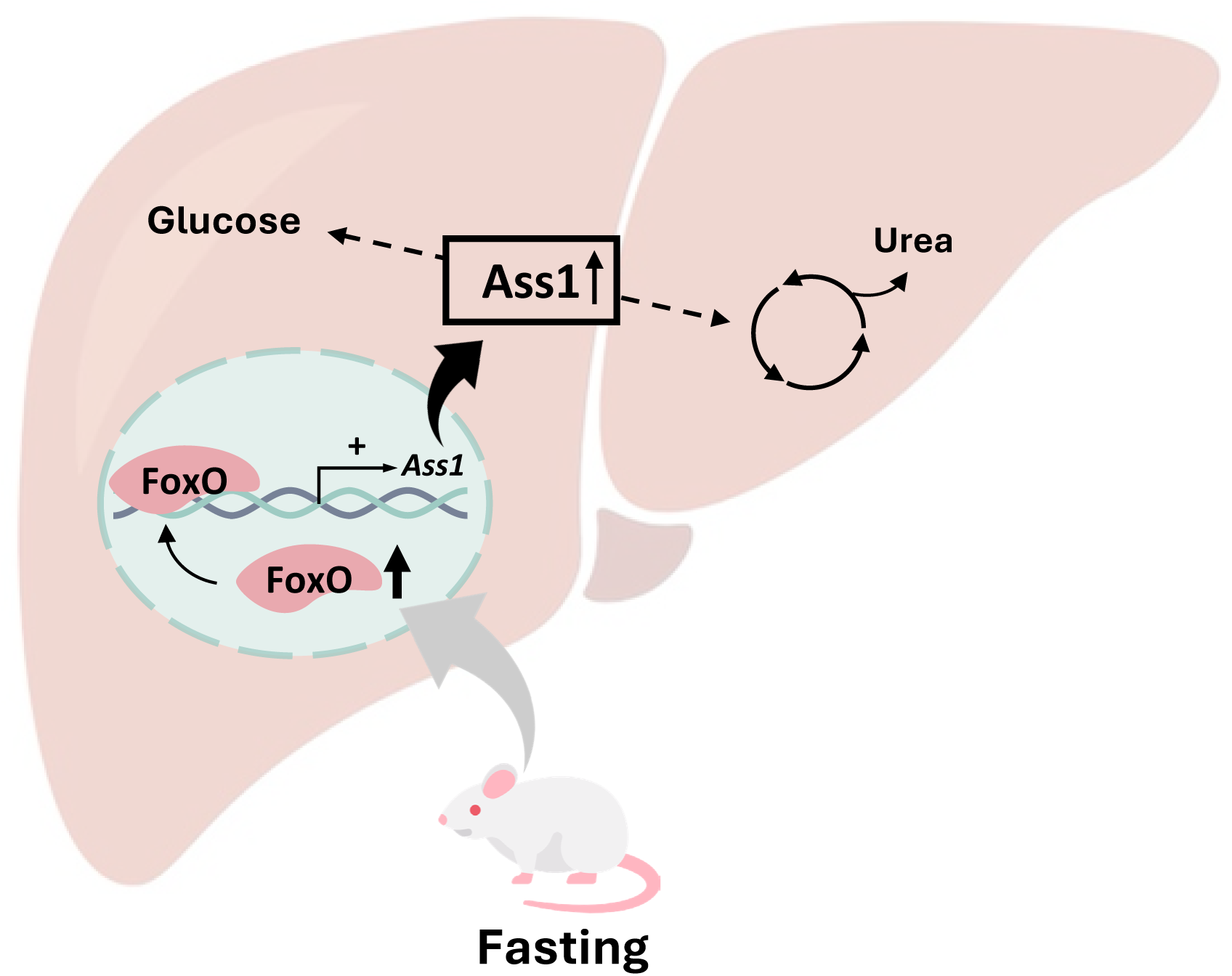

## Introduction

Far beyond their role as proteinogenic molecules, amino acids serve as pivotal agents in metabolic integration ^1–3^, with their catabolic byproducts significantly influencing lipid and carbohydrate metabolism ^4^. Given this essential function, amino acid breakdown is subjected to strict regulatory control ^5,6^. Acting as the central orchestrator of metabolic integration, the liver dynamically processes amino acids converting them into corresponding metabolites in response to the organism’s metabolic state and cellular demands ^5,7^.

The deamination of amino acids generates nitrogenous waste in the form of free ammonia, a toxic byproduct that poses a metabolic burden on the organism ^4,8^. To prevent nitrogen accumulation and maintain homeostasis, this process is tightly linked to the urea cycle, which efficiently detoxifies ammonia by converting it into urea for excretion ^9–13^. Also known as the ornithine cycle, the urea cycle is a liver-specific pathway, comprises five core enzymes: carbamoyl-phosphate synthetase 1 (CPS1), ornithine transcarbamylase (OTC), argininosuccinate synthetase (ASS1), argininosuccinate lyase (ASL), and arginase 1 (ARG1) ^14^. Notably, ASS1 and ASL, though classically defined as urea cycle enzymes, also serve a gluconeogenic function through fumarate production. This dual role is often overlooked in assessments of gluconeogenic rate control, particularly in the segment between pyruvate and phosphoenolpyruvate ^8^. Like other enzymes involved in amino acid catabolism, the expression of urea cycle genes is upregulated in response to increased substrate availability, such as during starvation or high dietary protein intake, thereby supporting enhanced protein degradation ^15–20^. It’s worth noting that the enzymatic activities of urea cycle enzymes correlate with their mRNA levels, suggesting that their regulation occurs predominantly at the pre-translational level ^21^.

Starvation represents a critical challenge that requires tight metabolic adaptation ^7^. During starvation, enhanced proteolysis supplies amino acids whose carbon skeletons are redirected toward gluconeogenesis ^22–24^. When amino acids are required for energy, their catabolism is initiated by the removal of the α-amino group via transamination and oxidative deamination ^4,5^. The resulting carbon skeletons then enter central metabolic pathways, either being oxidized through the TCA cycle to generate ATP or funneled into gluconeogenesis for glucose synthesis ^7,25^. This reliance on gluconeogenesis brings it into several functional intersections with the urea cycle. First, upregulation of the urea cycle facilitates amino acid-based gluconeogenesis by removing amino groups from amino acids, thereby freeing their carbon skeletons for catabolic use ^23,24^. In addition, fumarate, which is produced from aspartate via the sequential actions of ASS1 and ASL within the urea cycle, plays a pivotal role in glucose synthesis, as it is converted to malate and subsequently oxidized to oxaloacetate, a key gluconeogenic precursor ^7,8,10,24,26^.

Over a century ago, Graham Lusk demonstrated the influence of amino acids on glucose metabolism through the D/N ratio ^27^, highlighting the longstanding interest in nutrient-driven metabolic control. Despite this, the mechanisms that coordinate nitrogen disposal with glucose production, particularly the transcriptional regulation of urea cycle enzymes, remain underexplored. This gap persists even though the urea cycle was characterized as early as 1932 by Hans Krebs and colleagues ^28^, several years before the discovery of the tricarboxylic acid (TCA) cycle in 1937 ^29^, whose transcriptional control is far better understood. Thus far, OTC remains the only urea cycle enzyme known to be directly regulated at the transcriptional level by Kruppel-like factor 15 (KLF15) ^30^, and the wider regulatory network responsible for coordinated urea cycle induction during nutrient deprivation has yet to be elucidated.

KLF15, a transcription factor highly expressed in metabolically active tissues such as the liver and kidneys ^31,32^, contributes to the circadian regulation of nitrogen metabolism ^33^, with *Klf15*-deficient mice exhibiting disrupted circadian rhythms of amino acid and urea levels, highlighting its essential role in temporal metabolic control^34^. Expanding on its known function in amino acid metabolism, our findings have revealed that KLF15’s role in amino acid metabolism is more extensive than previously recognized. Specifically, under conditions of high-protein diet-induced metabolic changes in the liver, KLF15-dependent and independent pathways of amino acid catabolism can be distinguished. KLF15 directly influences the metabolism of 11 of the 20 standard amino acids (A, P, Q, M, I, V, L, K, W, F, and Y), underscoring its broad and critical function in amino acid homeostasis ^35^. Additionally, we identified KLF15 as a key regulator of lipid metabolism pathways ^36^, in addition to its established roles in amino acid catabolism ^30^ and glucose metabolism ^37^, thereby integrating multiple nutrient-sensing signals to coordinate metabolic responses.

In line with the integrating role of KLF15, our previous work showed that the hepatic Forkhead box O (FoxO)-KLF15 axis serves as a key conductor in the metabolic network directing the flow of macronutrients between starvation and overnutrition by linking amino acid-driven gluconeogenesis and glucose-driven lipogenesis under the influence of insulin ^38^. FoxO transcription factors, well-recognized for their conserved role in promoting health span and longevity across diverse species ^39,40^, play a crucial role in coordinating energy metabolism ^41^. They regulate glucose homeostasis by stimulating both gluconeogenesis and glycogenolysis ^42–45^, while also modulating lipid metabolism through upregulation of genes involved in fatty acid oxidation ^46^. This multifaceted control positions FoxO transcription factors as key metabolic regulators bridging nutrient sensing and adaptive responses. Yet, regulatory mechanisms are more complex. In response to high-protein nutritional states, we recently identified hepatic *Ass1* as a KLF15-independent gene under the direct control of FoxO transcription factors, highlighting an additional layer of transcriptional regulation in amino acid metabolism that operates beyond the KLF15 axis ^47^.

In this study, we investigated the role of FoxO transcription factors in coupling amino acid catabolism to glucose production during starvation. Specifically, we focused on the regulation of *Ass1*, a key urea cycle gene, as a direct FoxO target, and explored how this regulatory interaction may functionally link ureagenesis and gluconeogenesis. By clarifying this connection, we aim to uncover a transcriptional mechanism by which the liver aligns nitrogen disposal with energy production during starvation.

## Results

### FoxO transcription factors regulate the metabolism of amino acids involved in the urea cycle independently of Klf15 in the liver during fasting

In earlier work we have successfully shown that during fasting hepatic FoxO transcription factors accelerate amino acid breakdown and suppress lipogenesis all through the positive regulation of *Klf15* ^38^. Nevertheless, the possibility that FoxO transcription factors directly regulate other amino acid metabolic pathways remains a question worthy of investigation. In order to elucidate the individual role of each transcription factor, we designed a four-group mouse experiment; wild-type and *Klf15* knockout mice each treated with adenovirus mediated *LacZ* RNAi (Ad-*LacZ*i) as a control and *FoxO1* and *FoxO3a* RNAi (Ad-*FoxO1,3a*i), which specifically targets both *FoxO1* and *FoxO3a* ^38^. Blood glucose levels and plasma and liver amino acid profiles were compared across the four groups in the fasted state. *Klf15* knockout mice showed lower blood glucose levels compared to the *Klf15* wild-type mice (Figure 1A). Additionally, *FoxO* knockdown groups in both wild-type and *Klf15* knockout mice showed a significant decrease in blood glucose levels, with the *Klf15* knockout- *FoxO* knockdown group exhibiting the lowest fasting blood glucose levels of all, likely due to the combined effect of reduced *Klf15* and *FoxO* expressions. These data are consistent with previous reports ^30,48^. Regarding amino acid profile, two of the amino acids related to the urea cycle (Figure 1B) were notably affected. Specifically, plasma arginine levels (Figure 1C) were significantly decreased with *FoxO* knockdown in both *Klf15* wild-type and knockout groups while *Klf15* knockout had no effect on its levels. By contrast, plasma ornithine levels (Figure 1D) showed a significant increase in the same groups, also no significant effect by *Klf15* knockout. Other urea cycle-related amino acids and metabolites, aspartate (Figure 1E), citrulline (F), urea (G), and ammonia (H) showed no significant change caused by *FoxO* knockdown or *Klf15* knockout. The complete plasma amino acid profile is shown in Figure S2. Branched chain amino acids (BCAA) leucine (Figure S2A), isoleucine (B) and valine (C), along with tyrosine (D), and proline (E) showed a significant increase in their plasma levels due to *Klf15* knockout, whereas glutamine (F) and glycine (G) showed a significant decrease. These data are consistent with previous reports ^30,33^. It is worth mentioning that serine (Figure S2I) and threonine (J) showed increased plasma levels with *FoxO* knockdown in both *Klf15* wild-type and knockout groups. To assess whether these metabolic changes are concordant across compartments, we measured hepatic amino acid levels in the previously analyzed mice. As shown in Figure 1I-N, urea cycle-related metabolite levels were consistent between liver and plasma under our experimental conditions. The unchanged plasma levels of urea are not indicative of maintained nitrogen homeostasis in plasma but rather reflects the presence of multiple feedback and compensatory regulatory loops that might control urea levels in many tissues ^49^. To ensure reproducibility of results, we analyzed both plasma and hepatic amino acids in ICR mice with *FoxO* knockdown in the fasted state. As Figure 1O, U, P and V show, and consistent with the previous results, arginine levels were significantly decreased, and ornithine levels were significantly increased in both plasma and liver. On the other hand, plasma and hepatic levels of aspartate (Figure 1Q, W), citrulline (R, X), urea (S, Y), and ammonia (T, Z) showed no significant difference with *FoxO* knockdown. Additional data on the full hepatic amino acid profile during fasting, including comparisons with the ad libitum-fed state, are provided in Figure S3.

**Figure 1.**
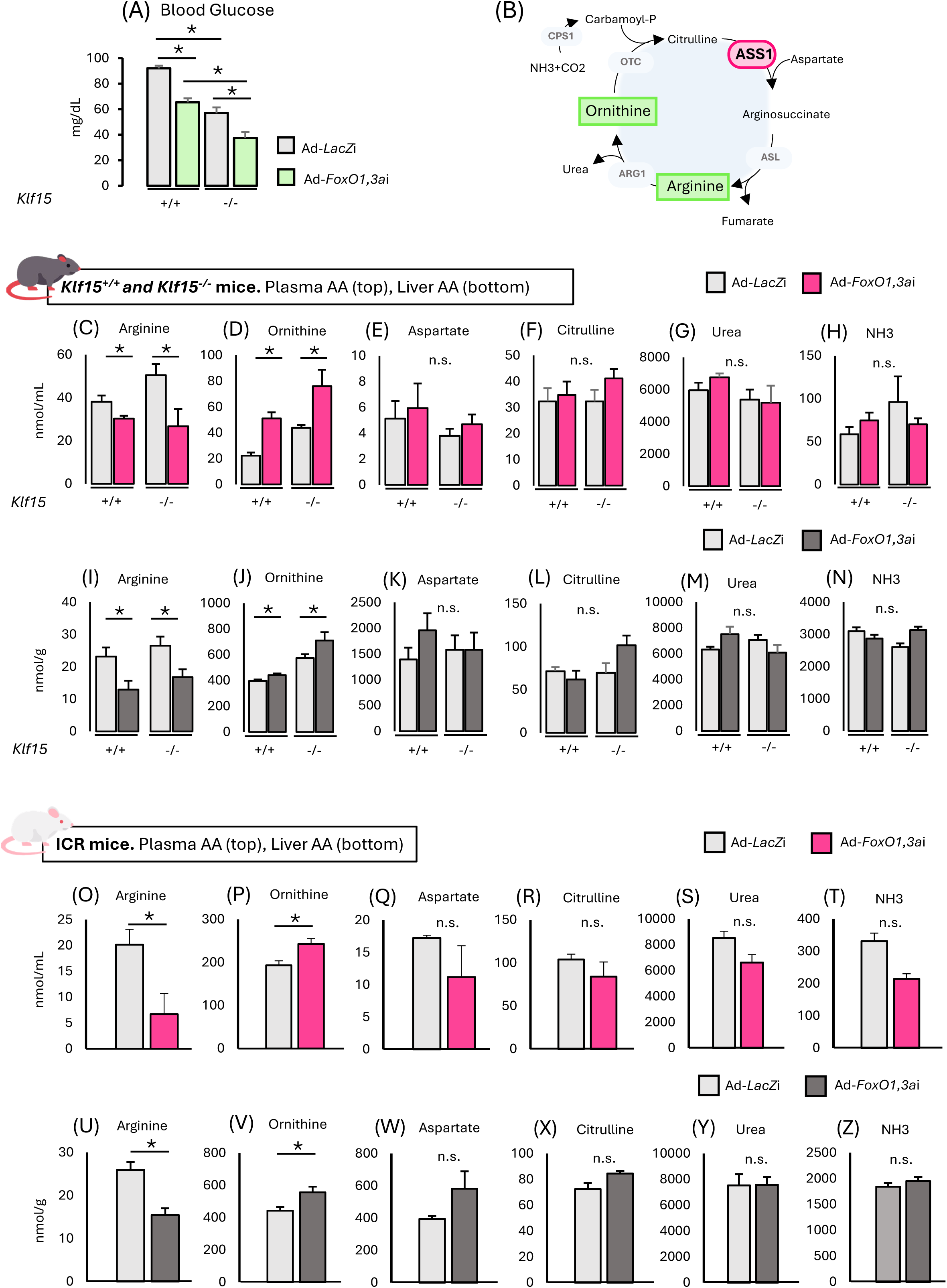
FoxO transcription factors regulate the metabolism of amino acids involved in the urea cycle independently of Klf15 in the liver during fasting. (**A**) Blood glucose levels in *Klf15* wild-type and knockout C57BL/6J mice with hepatic *FoxO1* and *FoxO3a* knockdown (n = 4 per group). (**B**) Schematic representation of urea cycle enzymes and related amino acids. (**C-H**) Plasma levels of urea cycle related amino acids, urea, and ammonia (NH3) in *Klf15* wild-type and knockout C57BL/6J mice with hepatic *FoxO1* and *FoxO3a* knockdown (n = 4 per group). (**I-N**) Hepatic levels of urea cycle related amino acids, urea, and ammonia (NH3) in *Klf15* wild-type and knockout C57BL/6J mice with hepatic *FoxO1* and *FoxO3a* knockdown (n = 4 per group). (**O-T**) Plasma levels of urea cycle related amino acids, urea, and ammonia (NH3) in ICR mice with hepatic *FoxO1* and *FoxO3a* knockdown (n = 5 per group). (**U-Z**) Hepatic levels of urea cycle related amino acids, urea, and ammonia (NH3) in ICR mice with hepatic *FoxO1* and *FoxO3a* knockdown (n = 5 per group). Plasma and hepatic amino acid levels were measured in ICR mice to ensure reproducibility of results. *FoxO1* and *FoxO3a* knockdown were performed using adenovirus mediated RNAi (Ad-*FoxO1,3a*i). All groups of mice were sacrificed at the same period in the light cycle and blood and liver samples were collected in the fasted state following 24-h fasting. All values are presented as the mean with error bars representing the SEM. Datasets were assessed by Student’s *t*-test for unpaired samples. The differences were considered to be significant if *P* < 0.05 (\**P* < 0.05 and \*\**P* <0.01).

### Hepatic FoxO transcription factors control the urea cycle through *Ass1* during fasting

To have a better understanding of the specific contribution of FoxO transcription factors-mediated amino acid metabolism related to the urea cycle, and building on our previous findings that FoxO transcription factors regulate argininosuccinate synthase 1 (*Ass1*) under high-protein diet conditions ^47^, we evaluated its gene expression levels in *Klf15* wild-type and knockout mice treated with both Ad-*LacZ*i and Ad-*FoxO1,3a*i during fasting. As shown in Figure 2A, and in line with the previously shown amino acids plasma and hepatic levels, *Ass1* gene expression was significantly decreased with *FoxO* knockdown independently of *Klf15* knockout. *Pck1* (Figure 2B) and *G6pc* (2C), known targets of FoxO transcription factors, also showed decreased gene expression with *FoxO* knockdown again with no difference between *Klf15* wild-type and knockout groups. *Arg1* (Figure 2D) expression showed no change across *FoxO* knockdown and *Klf15* knockout groups, and its expression level was used as a negative control. The gene expression levels were also assessed in ICR mice treated with both Ad-*LacZ*i and Ad-*FoxO1,3a*i to ensure reproducibility of results. In accordance with the abovementioned data, *Ass1*, *Pck1* and *G6pc* gene expression showed a significant decrease with lower *FoxO* gene expression (Figure 2G-I). The adenovirus-mediated overexpression of dominant-negative FoxO1 (Ad-FoxODN) ^38^ exhibited essentially the same effects on expression levels of these genes to exclude the possibility of artificial effects (Figure 2M-R). Figure 2E, F, K and L show the knockdown efficiency of both *FoxO1* and *FoxO3a* using Ad-*FoxO1,3a*i, and Figure 2Q shows FoxO1 protein over-expression using Ad-FoxODN. Additional comparisons of gluconeogenic and urea-cycle gene expression, together with Ass1 protein levels in total liver lysates from fasted and ad libitum-fed mice, are shown in Figure S4.

**Figure 2.**
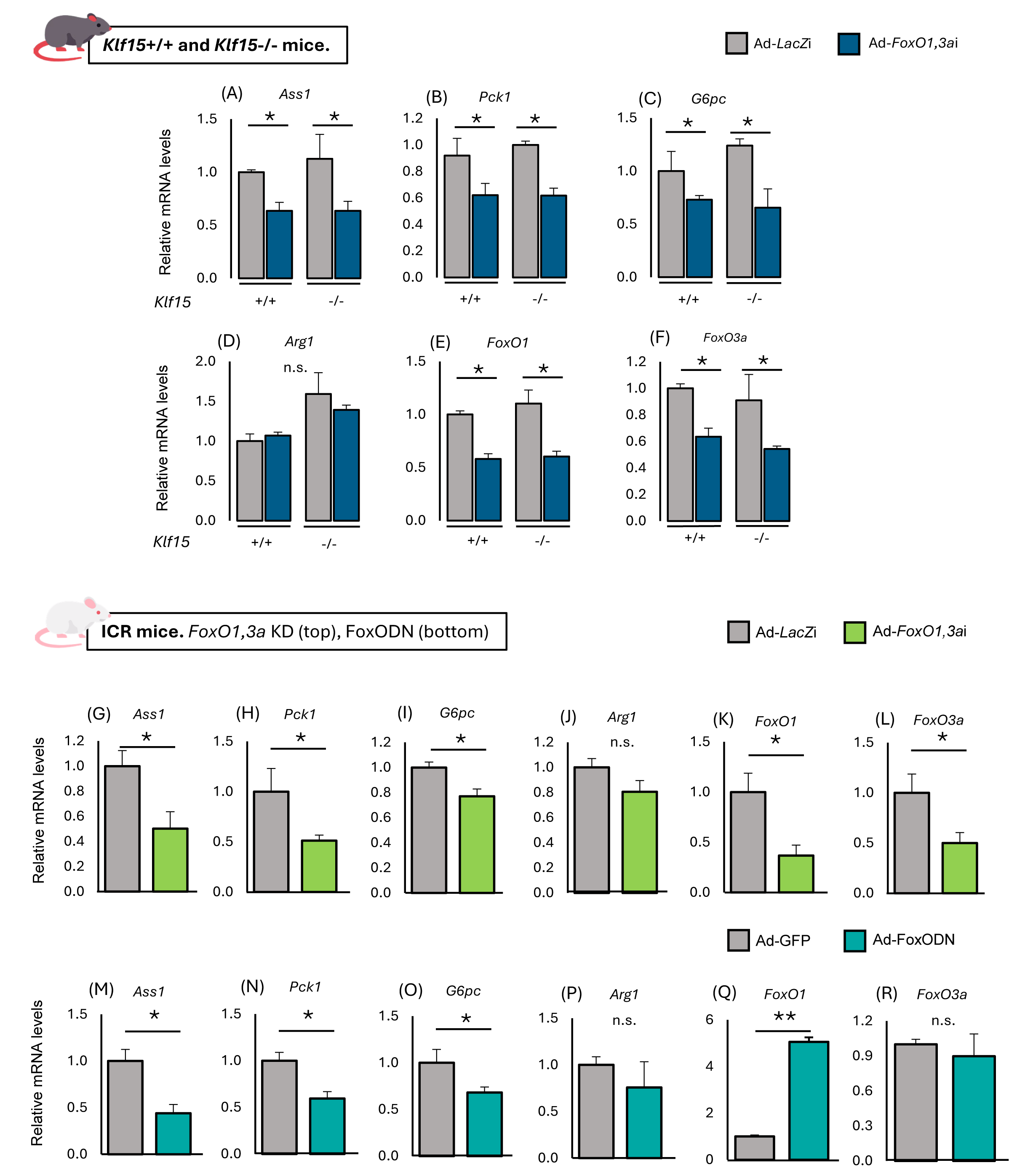
Hepatic FoxO transcription factors control the urea cycle through *Ass1* during fasting. Q-RT PCR analysis of liver RNA samples. (**A**-**F**) Relative gene expression in *Klf15* wild-type and knockout C57BL/6J mice with hepatic *FoxO1* and *FoxO3a* knockdown (n = 4 per group). (**G**-**L**) Relative gene expression in ICR mice with hepatic *FoxO1* and *FoxO3a* knockdown (n = 5 per group). (**M**-**R**) Relative gene expression in ICR mice with FoxO1 dominant negative overexpression (n = 4 per group). Relative gene expression was analyzed in ICR mice with *FoxO1,3a* knockdown to ensure reproducibility of results. Relative gene expression was analyzed with FoxO1DN to exclude the possibility of artificial effects. As the correction of the gene expression level for each sample *Cyclophilin A* was used in (**A**-**L**). *Gapdh* was used in (**M**-**R**). *FoxO1* and *FoxO3a* knockdown were performed using adenovirus mediated RNAi (Ad-*FoxO1,3a*i). FoxO1 dominant negative protein over-expression was performed using protein expressing adenovirus (Ad-FoxODN). All groups of mice were sacrificed at the same period in the light cycle and liver samples were collected in the in the fasted state following 24-h fasting. All values are presented as the mean with error bars representing the SEM. Datasets were assessed by Student’s *t*-test for unpaired samples. The differences were considered to be significant if *P* < 0.05 (\**P* < 0.05 and \*\**P* <0.01).

### Metabolic phenotypes analogous to *FoxO* knockdown were observed following *Ass1* knockdown during fasting

Hepatic *Ass1* was silenced with two adenoviral RNAi constructs (*Ass1*i-1 and *Ass1*i-2) ^47^ to assess its potential role in mediating the regulation of urea cycle-related amino acid metabolism by FoxO transcription factors. Blood glucose levels and plasma and liver amino acid profiles were compared across the groups in the fasted state. Both *Ass1* knockdown groups showed lower blood glucose levels compared to Ad-*LacZ*i group (Figure 3A) which mirrors the phenotype caused by *FoxO* knockdown (Figure 1A). Amino acid measurement in the plasma of ICR mice during fasting revealed that *Ass1* knockdown phenocopied the effect of *FoxO* knockdown (Figure 3B-G). Arginine levels were decreased (B) and ornithine levels (C) were increased, whereas aspartate (D), citrulline (E), urea (F), and ammonia (G) levels showed no significant difference. Upon analyzing total liver lysate of ICR mice with *Ass1* knockdown (Figure 3H), we detected Ass1 successful knockdown in *Ass1*i-1 and *Ass1*i-2 treated groups during fasting. *Ass1* RNAi induced the knockdown of endogenous *Ass1* to levels comparable with the downregulation of *Ass1* upon *FoxO* knockdown (Figure 3I). At the same time, *FoxO1*, *FoxO3a*, and other urea cycle genes showed no differences in *Ass1*i groups (Figure 3J-Q). The two small hairpin RNAs (shRNAs) exerted essentially the same effects on amino acid levels and the expression levels of other urea cycle genes, excluding the possibility of off-target effects. Collectively, our findings suggest that *Ass1* is a critical direct target of the FoxO transcription factors in the regulation of the urea cycle-related amino acid metabolism during fasting.

**Figure 3.**
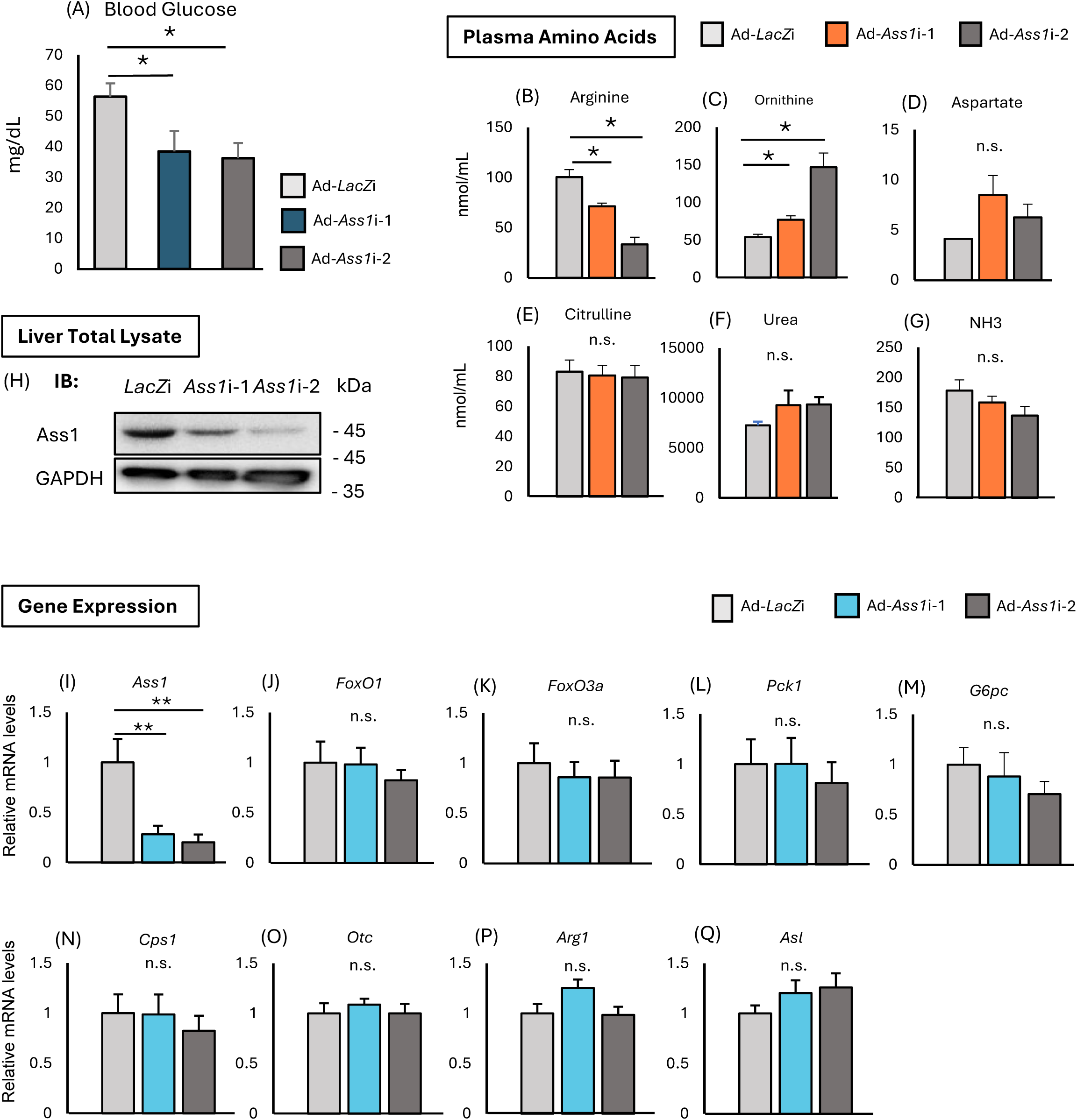
Metabolic phenotypes analogous to *FoxO* knockdown were observed following *Ass1* knockdown during fasting. (**A**) Blood glucose levels in ICR mice with hepatic *Ass1* knockdown (n = 4 per group). (**B**-**G**) Plasma levels of urea cycle related amino acids, urea, and ammonia (NH3) in ICR mice with hepatic *Ass1* knockdown (n = 4 per group). (**H**) Immunoblot analysis of Ass1 protein using liver total lysate from mice in the fasted state (Samples were pooled from 3 mice). GAPDH was detected as an internal loading control. (**I**-**Q**) Relative gene expression in ICR mice with hepatic *Ass1* knockdown (n = 4 per group). *Ass1* knockdown was performed using adenovirus mediated RNAi (Ad-*Ass1*i-1, Ad-*Ass1*i-2). As the correction of the gene expression level for each sample *Cyclophilin A* was used. All groups of mice were sacrificed at the same period in the light cycle and blood and liver samples were collected in the fasted state following 24-h fasting. All values are presented as the mean with error bars representing the SEM. Datasets were assessed by Student’s *t*-test for unpaired samples. The differences were considered to be significant if *P* < 0.05 (\**P* < 0.05 and \*\**P* <0.01).

### Identification of FoxO transcription factors binding region upstream of *Ass1* gene

To determine FoxO transcription factors binding sites in the *Ass1* enhancer region (Fig S1), we performed enhancer analysis using various *Ass1*-enhancer-luc constructs in HepG2 cells. Possible FoxO transcription factors binding sites in the enhancer region upstream of the *Ass1* gene has been identified using ChIP-Atlas (Figure 4A-Top). In order to locate the binding site, the mentioned area was subdivided into smaller fragments and cloned, along with the full-length fragment (Figure 4A-Bottom), into luciferase reporter plasmid to be studied separately through evaluating the luciferase activity in HepG2 cells. As the data show, both full-length and fragment 1 (Figure 4B) and subdivision A of fragment 1 (Figure 4C) showed significant luciferase activity to both FoxO1 and FoxO3a expression plasmids compared to other fragments and to the control.

**Figure 4.**
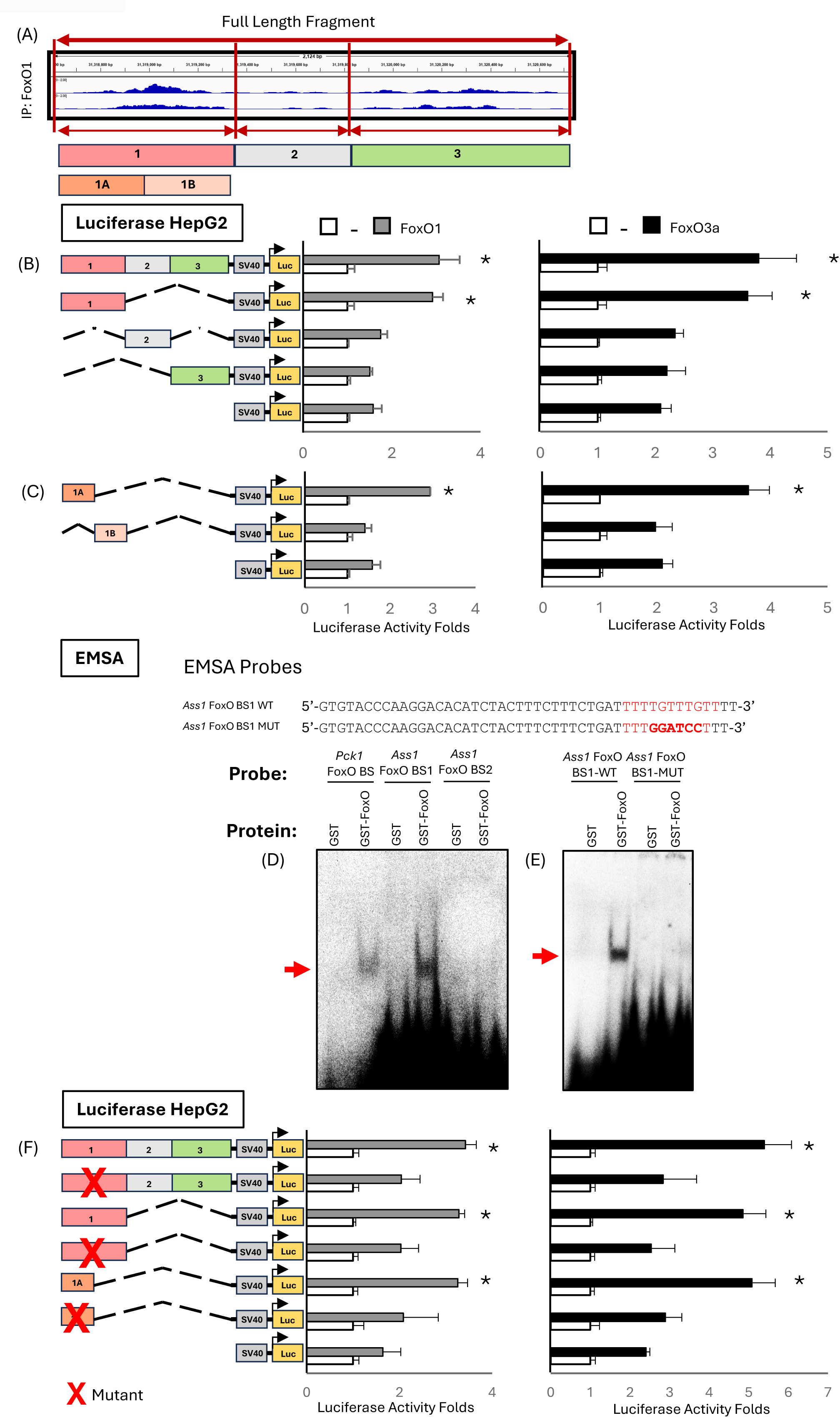
Identification of FoxO transcription factors binding region upstream of *Ass1* gene. (**A**) Top, the results of ChIP-seq with anti-FoxO1 antibody in livers of unfed mice obtained from ChIP-Atlas (ID: GSM3381273); bottom, simplified representation of the structure of the full fragment, fragment 1, fragment 2, fragment 3, and the subdivisions of fragment 1 i.e. small fragment 1A and small fragment 1B. (**B** and **C**) Enhancer analysis of FoxO1 and FoxO3a on *Ass1* enhancer and its fragments. FoxO1 and FoxO3a expression plasmids were co-transfected with the indicated *Ass1* enhancer firefly luciferase reporter plasmids in HepG2 cells. (**D** and **E**) Electrophoretic mobility shift assay (EMSA) using radiolabeled probe for (**D**) the two JASPAR predicted FoxO binding sites (Figure S5) in the smallest fragment 1A from the *Ass1* enhancer and FoxO binding site in the promoter of *Pck1* as a positive control; (**E**) the wild-type (WT) and mutant (Mut) predicted FoxO binding site1 incubated with GST and GST-FoxO recombinant proteins. BS, binding site; WT, wild-type; Mut, mutant. (**F**) Enhancer analysis of FoxO1 and FoxO3a on mutant *Ass1* enhancer and its fragments. FoxO1 and FoxO3a expression plasmids were co-transfected with the indicated *Ass1* enhancer firefly luciferase reporter plasmids in HepG2 cells. All values are presented as the mean with error bars representing the SEM. Datasets were assessed by Student’s *t*-test for unpaired samples. The differences were considered to be significant if *P* < 0.05 (\**P* < 0.05 and \*\**P* <0.01).

To support these findings, an EMSA assay was performed. For that, two radiolabeled probes for the two high score binding sites in fragment 1A based on JASPAR analysis results were designed (Figure S5A-D) and checked against FoxO binding site on *Pck1* promoter as a positive control. EMSA results showed that the recombinant FoxO protein binds to the DNA radiolabeled probe of binding site number 1 (*Ass1* FoxO BS1) (Figure 4D). Then as a next step we designed a probe for mutated *Ass1* FoxO BS1 and no band was observed (Figure 4F).

Consistent with these findings, when we mutated this predicted binding site in the full-length fragment, fragment 1 and fragment 1A, the activation of the luciferase gene by both FoxO1 and FoxO3a expression plasmids in HepG2 cells was completely abolished as shown in Figure 4F. Collectively, these findings identify the specific FoxO-binding site essential for enhancer-mediated activation

### The identified region is required for hepatic *Ass1* response to fasting

To validate these findings, we assessed the effect of fasting on transcriptional activity *in vivo* by engineering the smallest fragment containing the identified FoxO binding site (Figure 4) both the wild-type and the mutated versions into a luciferase reporter plasmid connected to *Ass1* native promoter (Figure 5A) and delivered adenovirally to the liver. The transcriptional activity was assessed by measuring luciferase activity with an IVIS imaging system ^36,50–52^ before and after fasting. As shown in Figure 5B-F the reporter activity of the wild-type FoxO binding site construct significantly increased in the fasted state compared to the fed state (5C, D) and compared to the mutated version (5C, E). While the response of the mutated FoxO binding site construct was significantly attenuated (5C, E, F) suggesting that this site is necessary for the activation of *Ass1*.

**Figure 5.**
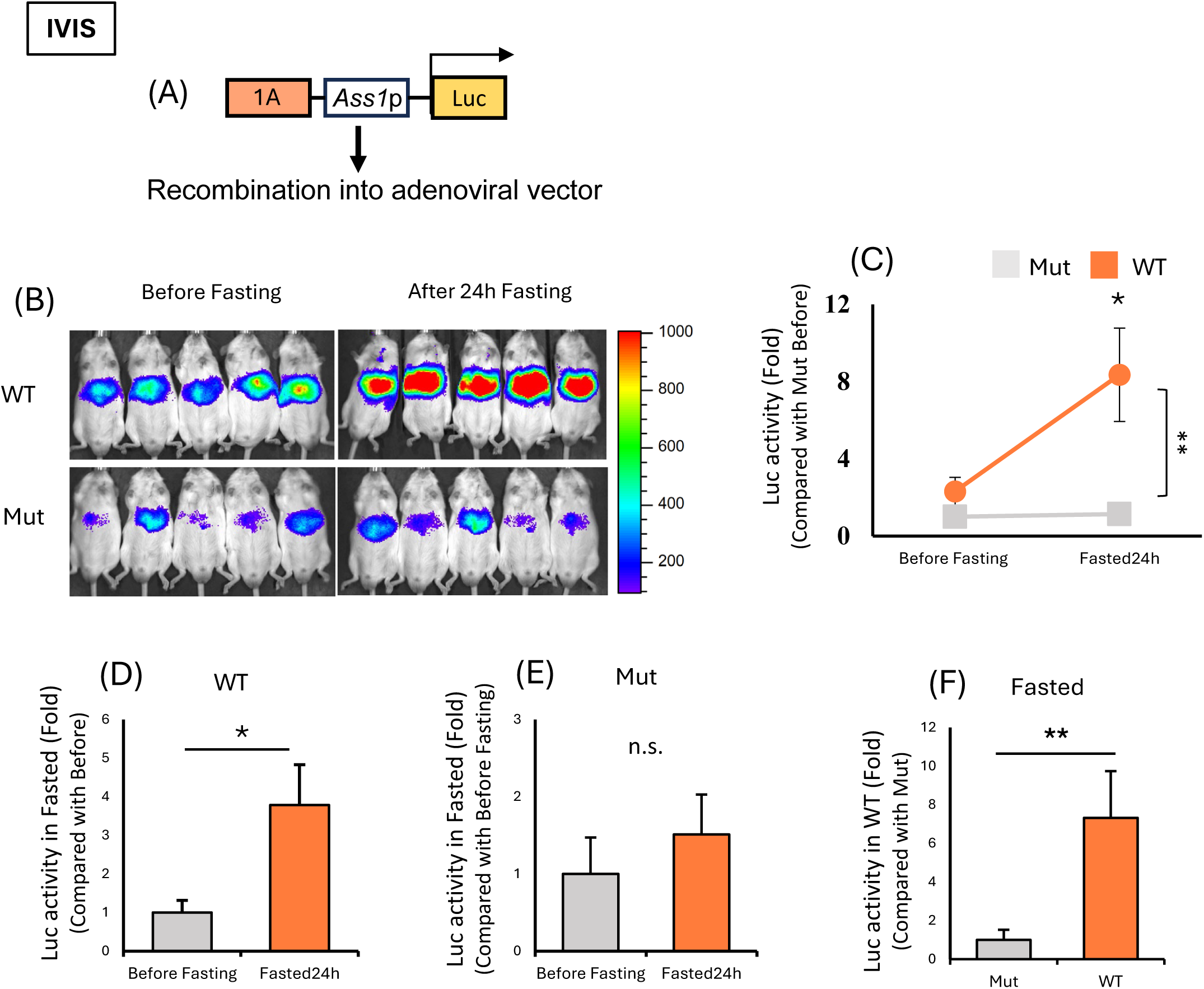
The identified region is required for hepatic *Ass1* response to fasting. (**A**) The structure of the Ad-Luc showing the identified FoxO binding site in the enhancer region of *Ass1* linked to the Luc reporter using *Ass1* native promoter. (**B**-**F**) In vivo Ad-Luc enhancer analyses using Ad-FoxOBS-*Ass1*Promoter-Luc both wild-type and mutant. Representative images (**B**) and hepatic luciferase activities (**C**-**F**) of mice injected with Ad-FoxOBS-*Ass1*Promoter-Luc both wild-type and mutant are shown (n = 5). (**D**) Enhancer activity in the liver for wild-type binding site after 24-h fasting is expressed relative to activity before fasting, to adjust for mouse-to-mouse differences in enhancer expression in Ad-treated mice. (**E**) Enhancer activity in the liver for mutant binding site after 24-h fasting is expressed relative to activity before fasting, to adjust for mouse-to-mouse differences in enhancer expression in Ad-treated mice. (**F**) Enhancer activity in the liver after 24-h fasting for wild-type binding site is expressed relative to activity of mutant binding site, to adjust for mouse-to-mouse differences in enhancer expression in Ad-treated mice. BS, binding site; WT, wild-type; Mut, mutant. Data were assessed using the paired two-tailed Student’s *t*-test. The differences were considered to be significant if *P* < 0.05 (\*\**P* < 0.01). Error bars mean SEM.

### FoxO transcription factors bind to hepatic *Ass1* enhancer during fasting

To have a deeper insight into the regulatory mechanism of FoxO transcription factors over *Ass1* gene expression in vivo, we employed chromatin immunoprecipitation (ChIP) assay to evaluate the binding ability of FoxO transcription factors to *Ass1* enhancer during the fasted state compared to the ad libitum-fed state using both anti-FoxO1 and anti-FoxO3a antibodies against anti-IgG antibody. As Figures 6A,E depicted, FoxO1 and FoxO3a binding occupancy to the identified FoxO binding site on the *Ass1* enhancer region (Figure 4) was significantly increased during fasting compared with the ad libitum-fed state. *Pck1* (6B,F) and *G6pc* (6C,G) promoter FoxO transcription factor binding site were used as a positive control and showed FoxO transcription factor occupancy on the corresponding promoter. *Ass1* gene body (5D,H) was used as negative controls. Next, upon examining hepatic nuclear FoxO1/FoxO3a protein levels (Figure 6I) we show how nuclear FoxO1 and FoxO3a protein levels increase during fasting compared to ad libitum-fed. Taken all together, these findings suggest that nuclear FoxO transcription factor availability during fasting directly stimulate *Ass1* transcription in the liver.

**Figure 6.**
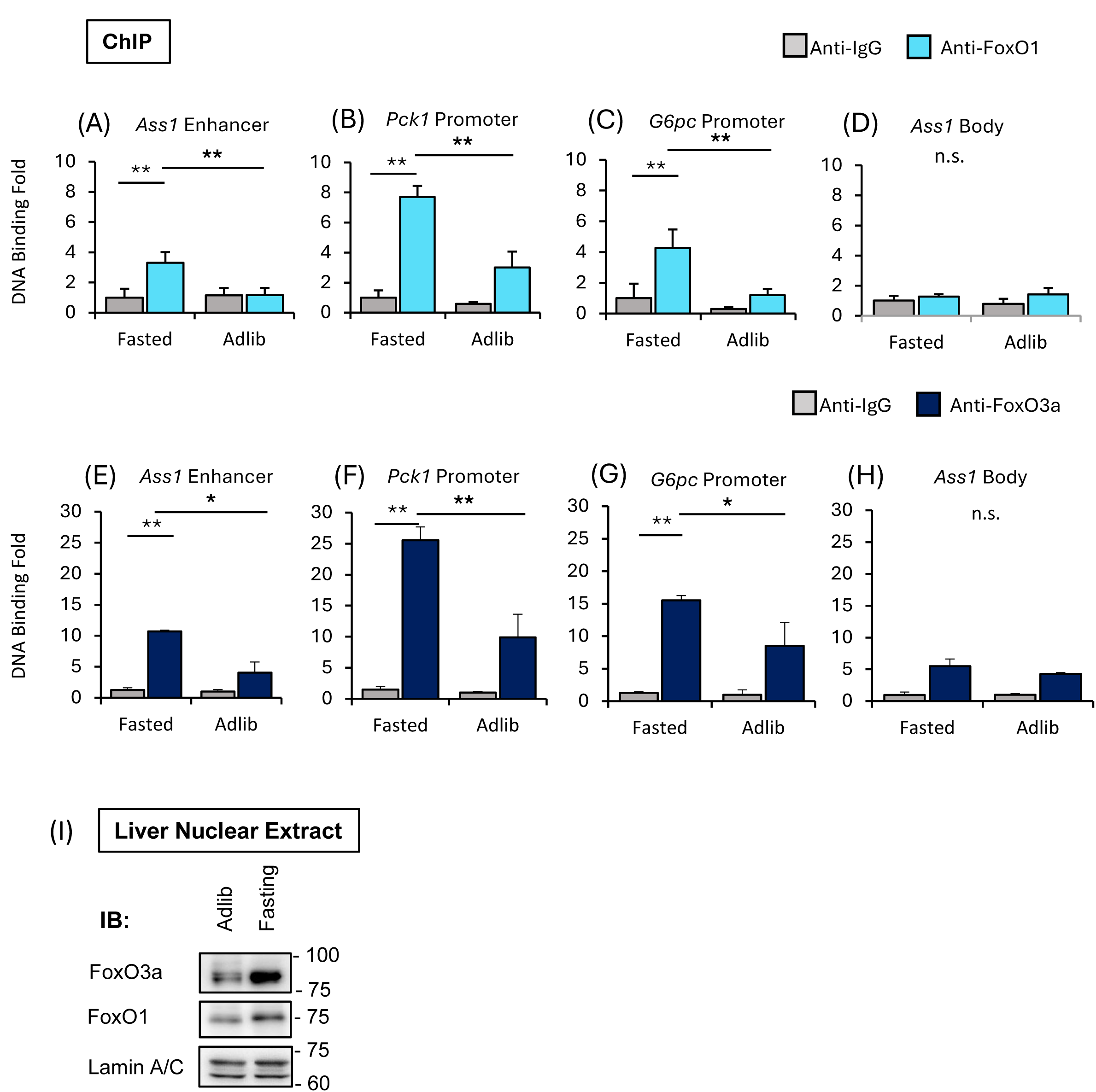
FoxO transcription factors bind to hepatic *Ass1* enhancer during fasting. (**A**-**D**) Chromatin immunoprecipitation (ChIP) assay to determine the interaction of hepatic FoxO proteins with *Ass1* DNA enhancer region using anti-FoxO1 antibody and normal mouse anti-IgG antibody as a control. (**E**-**H**) Chromatin immunoprecipitation (ChIP) assay to determine the interaction of hepatic FoxO proteins with *Ass1* DNA enhancer region using anti-FoxO3a antibody and normal mouse anti-IgG antibody as a control. (**A**, **E**) FoxO proteins binding to *Ass1* enhancer in liver detected by using primer set designed around the identified FoxO binding site in Figure 4. (**B**, **F**) *Pck1* promoter FoxO transcription factor binding site and (**C**, **G**) *G6pc* promoter FoxO transcription factor binding site used as a positive control. (**D**, **H**) Used as a negative control. (**I**) Immunoblot analysis of FoxO1 and FoxO3a proteins using liver nuclear extracts from mice in ad libitum-fed and fasted state (Samples were pooled from 3 mice). Lamin A/C was detected as an internal loading control for nuclear protein. All groups of mice were sacrificed in the light cycle and liver samples were collected in the fasted state following 24-h fasting or ad libitum-fed as indicated. Data were assessed using the unpaired two-tailed Student’s *t*-test. The differences were considered to be significant if *P* < 0.05 (\*\**P* < 0.01). Error bars mean SEM.

## Discussion

In this study, we identified a novel role for FoxO transcription factors as direct regulators of hepatic *Ass1*, a key urea cycle gene. This regulation occurs independently of Klf15 and orchestrates changes in urea cycle-related amino acid levels during starvation. By investigating this FoxO-*Ass1* interaction, we revealed a transcriptional mechanism that may functionally couple ureagenesis and gluconeogenesis, aligning nitrogen disposal with the energy production needs of the liver during nutrient deprivation.

To identify FoxO transcription factors as upstream regulators of *Klf15*, our group previously employed a comprehensive Transcription Factor Expression Library (TFEL), which encompasses nearly all transcription factors encoded in the mouse genome ^53^. Through this approach, we demonstrated that hepatic FoxO transcription factors control amino acid metabolism via the Klf15 pathway ^38^. Furthermore, we recently confirmed that hepatic *Ass1* is directly regulated by FoxO transcription factors in response to high-protein nutritional states, revealing a branch of transcriptional control in amino acid metabolism that functions independently of the Klf15 axis ^47^.

To adapt to fluctuations in caloric intake, organisms adjust the balance between carbohydrate and amino acid utilization as energy sources ^25^. When amino acids are catabolized to meet energy demands, the resulting amino nitrogen is primarily converted into urea and excreted, reflecting the body’s capacity to maintain nitrogen balance while utilizing the carbon skeletons for energy ^14^. The upregulation of the five urea cycle enzymes during starvation is well documented (Figure S4 A,G-K) ^17^. However, the transcriptional mechanisms that regulate this adaptive response have remained largely unresolved. To date, only *OTC* has been identified as a transcriptional target of KLF15 ^30^, leaving the broader regulatory landscape governing urea cycle activation during nutrient deprivation still poorly understood.

Our data revealed that reduced availability of FoxO transcription factors significantly affected the plasma and hepatic levels of arginine and ornithine (Figure 1). Interestingly, the concentrations of citrulline and aspartate, the two direct substrates of Ass1, as well as ammonia and urea, remained largely unchanged in both plasma and liver. Consistently, the expression of *Arg1*, which hydrolyzes arginine into urea and ornithine, also remained stable (Figure 2). This overall stability of the urea cycle, despite perturbations in individual components, can be attributed to the inherent robustness and adaptability of cellular metabolic networks. Such networks maintain homeostasis through redundant pathways, feedback regulation, and compensatory mechanisms that buffer against fluctuations in gene expression and environmental stressors ^54^. These characteristics, along with the nonlinear dynamics and the complexity of interactions among numerous metabolites, often obscure the identification of discrete regulatory nodes within metabolic circuits ^49^.

During starvation, when amino acid catabolism is elevated, the urea cycle is pushed to operate at full capacity. Under these conditions, ASS1 functions as the rate-limiting step, with its activity primarily regulated by hormonal and nutritional cues^55^. Glucocorticoids and glucagon are known to enhance ASS1 activity, while insulin suppresses it ^56^. Despite the central role of ASS1 in regulating ureagenesis, only a single transcription factor, Sp1, has been experimentally confirmed to bind its promoter region to date ^57^. In the present study, we identified FoxO transcription factors as novel direct regulators of hepatic *Ass1* expression during starvation, acting independently of Klf15 (Figure 2). We further demonstrated that FoxO-binding elements within the *Ass1* enhancer are essential for its transcriptional activation in response to nutrient deprivation (Figure 5).

There are two potential mechanisms by which FoxO binding to the *Ass1* enhancer may increase. One possibility is an increased amount of nuclear FoxO protein caused by reduced phosphorylation by the insulin/PI3K/PDK/AKT cascade during fasting (Figure 6I) ^58–60^. The other possibility is that increased FoxO binding to the *Ass1* enhancer is caused by elevated amino acid concentrations during fasting (Figure S3 I-L), as demonstrated in our previous study using a high protein diet ^47^.

The contribution of FoxO transcription factors to amino acid metabolism remains incompletely defined, despite their well-established roles in numerous other metabolic processes. For instance, liver-specific ablation of *FoxO1*, *FoxO3*, and *FoxO4* (L-FoxO1,3,4) has been shown to impair the fasting-induced expression of glucose-6-phosphatase and prevent the repression of glucokinase, highlighting their importance in glucose homeostasis during nutrient deprivation ^61^. However, elucidating their connection to *Ass1* regulation is complicated by compensatory mechanisms. In the available microarray dataset (GSE60527/GPL6096/6768261), *Ass1* expression levels remained unchanged (13.27 in both hepatocyte-specific FoxO1,3,4 knockout and littermate control), suggesting that long-term FoxO depletion may trigger compensatory transcriptional programs that obscure functional relationships. Such compensation is a well-documented phenomenon in gene knockout models, particularly involving transcription factors, and often leads to misinterpretation of regulatory mechanisms. Comparative studies have demonstrated that knockouts and knockdowns can produce markedly different phenotypes across species, including mice ^62,63^. For example, in mice lacking SREBP-1, hepatic SREBP-2 expression is upregulated to compensate for the genetic loss of SREBP-1 ^64^. In this context, knockdown strategies, by inducing acute and partial gene suppression, may provide a more accurate representation of direct regulatory effects, as they are less likely to provoke systemic compensatory adaptations. It is well established that gluconeogenesis and ureagenesis function cooperatively to support sustained amino acid catabolism in the liver, primarily by generating ATP and facilitating nitrogen disposal ^15^. In liver-specific *FoxO1* transgenic mice, previous studies have shown an upregulation of genes involved in amino acid catabolism and gluconeogenesis, highlighting the role of FoxO1 in coordinating these metabolic pathways in the liver ^4^. This coordination ensures efficient processing of amino acids, maintaining metabolic balance during nutrient deprivation. These findings underscore the need to investigate the role of FoxO transcription factors as potential integrators of these pathways through the regulation of amino acid metabolism. In this context, our data demonstrated that hepatic FoxO depletion leads to partial disruption of the urea cycle and a significant reduction in blood glucose levels (Figure 1). To determine whether Ass1 mediates the regulatory effect of FoxO on urea cycle-related amino acid metabolism, we performed hepatic *Ass1* knockdown. Interestingly, *Ass1* depletion produced a phenotype resembling that of *FoxO* knockdown: both resulted in partially impaired urea cycle function, evidenced by altered ornithine and arginine levels, and lowered blood glucose, even though *Pck1* or *G6pc* gene expression remained unchanged (Figure 3). These findings suggest that *Ass1* is a critical direct target of FoxO transcription factors, mediating their role in integrating ureagenesis and gluconeogenesis. Thus, Ass1 may represent a key regulatory node linking nitrogen disposal and glucose production during starvation.

The coupling between ureagenesis and gluconeogenesis is physiologically essential during nutrient deprivation, when amino acids serve as a primary energy source ^5,24^. This coordination allows nitrogen to be safely eliminated via the urea cycle, while the resulting carbon skeletons are directed toward glucose production to maintain systemic energy homeostasis ^10^. A key point of intersection between these two pathways occurs through the conversion of aspartate to fumarate, alongside the transformation of citrulline to arginine ^8^. Although oxaloacetate cannot directly traverse the mitochondrial membrane, it can reach the cytosol after transamination to aspartate. This aspartate is then reconverted to oxaloacetate via fumarate and malate, a process that requires ASS1 activity ^65^. Our data support the view that, in the liver, the dominant net route by which aspartate-derived carbon contributes to cytosolic oxaloacetate proceeds through the urea cycle ^8^. Aspartate entry via ASS1, followed by argininosuccinate cleavage to fumarate, enables the return of carbon skeletons to the malate-oxaloacetate pool. In contrast, the reversible aspartate aminotransferase reaction contributes little net oxaloacetate production under steady-state conditions, despite high bidirectional flux.

Clinical evidence from inborn errors of metabolism underscores the physiological coupling between the urea cycle and gluconeogenesis, particularly under fasting conditions. For instance, in pyruvate carboxylase deficiency, impaired conversion of pyruvate to oxaloacetate disrupts both the TCA cycle and gluconeogenesis, leading to fasting-induced hypoglycemia ^66^. Crucially, oxaloacetate is also required for generating aspartate, a key substrate in the urea cycle. Its deficiency reduces argininosuccinate synthesis, resulting in impaired ureagenesis and hyperammonemia highlighting how a primary gluconeogenic defect can secondarily impair nitrogen disposal ^67^. Conversely, in Citrin (AGC2) deficiency, defective mitochondrial export of aspartate and malate reduces cytosolic availability of both substrates, limiting urea cycle activity and simultaneously impairing gluconeogenesis ^68^. This dual impact leads to recurrent hyperammonemia, hypoglycemia, and citrullinemia, caused by insufficient aspartate for urea synthesis and diminished fumarate generation for oxaloacetate production and subsequently gluconeogenesis ^69^. Together, these disorders illustrate the bidirectional metabolic interdependence between ureagenesis and gluconeogenesis, particularly under conditions of nutrient deprivation.

In summary, our study identifies FoxOs as central coordinators of the urea cycle and gluconeogenesis through direct transcriptional control of Ass1. This FoxO-Ass1 pathway provides a mechanistic explanation for how amino acid availability is translated into coordinated nitrogen excretion and carbon reutilization. Thus, FoxOs ensure metabolic flexibility during fasting.

## Acknowledgements

We thank Prof. Mukesh Jain (Case Western Reserve University) for kindly providing us with the *Klf15* knockout mouse.

This work was supported by MEXT/JSPS KAKENHI Grant Numbers 23116006 (Grant-in-Aid for Scientific Research on Innovative Areas: Crosstalk of transcriptional control and energy pathways by hub metabolites), 23K24760 (Grant-in-Aid for Scientific Research (B)), and 24K22110 (Grant-in-Aid for Challenging Exploratory Research) (to N. Yahagi). This research was also supported by AMED under Grant Number JP23gm1710008 (AMED-CREST) and JP23rea522010 (Healthcare Social Implementation Infrastructure Development Project) (to N. Yahagi). It was also supported by MEXT/JSPS KAKENHI Grant Number 25K24300 (Research Activity Start-up) (to S. Karkoutly).

## Author contributions

S.K. and N.Y. conceived the experiments. S.K. performed the experiments under the guidance of Y.T. and analyzed the data together with N.Y. S.K. Y.T. and N.Y. co-wrote the paper. All authors discussed the results and commented on the manuscript.

## Methods

### Animals

Male ICR mice aged between 5 to 7 weeks were purchased from Japan SLC, Inc. (Shizuoka, Japan). The *Klf15*^-/-^ (*Klf15*KO) backcrossed into C57BL/6J strain mice was kindly gifted by Prof. Jain MK ^70^. Controls were C57BL/6J *Klf15*^+/+^ (wild-type) mice. The experiments using ICR were performed between 6 and 8 weeks of age, and the experiments using *Klf15*^+/+^ and *Klf15*^-/-^mice were performed at 5-6 months of age. The animals were housed in a temperature-controlled environment with a 12-hour light/12-hour dark cycle and provided free access to standard laboratory diet and water. After introducing the standard laboratory diet (Cat#MF; Oriental Yeast, Tokyo, Japan; consisted of 25.5% of energy from protein, 61.5% from carbohydrates, and 13% from fat), mice were fasted for 24 hours, and blood glucose levels were measured from tail vein blood using a handheld glucometer before sacrifice. Mice were sacrificed during the early light phase in either ad libitum-fed or fasted states. All animal procedures were performed following the protocol approved by the Tsukuba University and Jichi Medical University Animal Care and Use Committee. The experiments were repeated at least twice to correct the bias for each experimental environment.

### Amino acid measurement

Amino acid measurements were performed as previously described ^47^. In brief, for plasma amino acids measurement, blood samples were collected from vena cava under a three-type mixed anesthesia consisting of medetomidine, midazolam, and butorphanol. Protein components were precipitated in the collected plasma samples by mixing them with equal amounts of 3% ice-cold sulfosalicylic acid. After centrifugation the supernatant was collected. The sample pH was adjusted to pH 2-3 and then filtered using a centrifuge filter (Cat#UFC30HV00, Millipore, Merck KGaA, Darmstadt, Germany). The whole process was performed on ice. The amino acid composition was measured using Hitachi-Hightech (LA8080).

For liver amino acids measurement, mouse liver samples were homogenized in 3% ice-cold sulfosalicylic acid to precipitate proteins. Later steps are as explained in plasma amino acids measurement.

### RNA extraction and Quantitative reverse transcription PCR (Q-RT PCR)

Total RNA was extracted from 50-100 mg of mouse liver using Sepasol-RNA I Super G (Cat# 09379-55; Nacalai Tesque, Kyoto, Japan) according to the manufacturer’s instructions. The extracted RNA (500 ng) was reverse transcribed in a volume of 5 μL and converted to cDNA using the ReverTra Ace qPCR RT Master Mix (Cat# FSQ-201, TOYOBO, Osaka, Japan). Real-time PCR was performed using KAPA SYBR Fast qPCR Kit (Cat# KK4602, NIPPON Genetics, Tokyo, Japan) on a QuantStudio^TM^ 5 Real-Time PCR System (Thermo Fisher Scientific, Waltham, MA, USA) and quantified by the standard curve method with cDNA as the template. After amplification by PCR, samples containing the product with the correct Tm value were taken based on the melting curve plot for each sample. Primer sets are listed in Table S1. *Cyclophilin A* was used as an internal reference to correct gene expression level for each sample.

### Plasmid construction

For constructing *Ass1*-enhancer firefly luciferase reporter plasmid, a 2.1 kbp-region of the 5’-flanking sequence at 7 kbp upstream from transcriptional start site on the mouse *Ass1* gene were chosen as the enhancer sequence for our experiments. Figure S1 shows UCSC Genome Browser tracks for H3K4me1, H3K27ac, and H3K9ac. The estimated enhancer site and its smaller fragments were constructed by amplifying genomic DNA by PCR using PrimeSTAR GXL DNA Polymerase (Cat#R050A, TaKaRa Bio, Tokyo, Japan) and linked to luciferase reporter gene by infusion technique using In-Fusion HD Cloning Kit (Cat#071320, TaKaRa Bio, Tokyo, Japan) according to the manufacturer’s protocol. Mutated *Ass1*- enhancer firefly luciferase reporter plasmids were generated using PrimeSTAR Mutagenesis Basal Kit (Cat# R046A, TaKaRa Bio, Tokyo, Japan). Used primers are listed in Table S2.

### Cell Culture

HEK293 human embryonic kidney cells (RRID:CVCL_0045) and HepG2 human hepatoma cells (RRID: CVCL_0027) were distributed from ATCC (American Type Culture Collection, Manassas, VA, USA), and cultured in DMEM containing 25 mM glucose, 100 U/mL penicillin, and 100 μg/mL streptomycin sulfate supplemented with 10% FBS. All cell lines have been authenticated by our facility’s authentication protocol within the last 3 years, and all experiments were performed with mycoplasma-free cells.

### Luciferase assay

For luciferase assay as described previously ^38^ cells were seeded in 48-well plate to 20% confluency and transfected using Lipofectamine 3000 reagent (Cat# L3000-015, Thermo Fisher Scientific, Waltham, MA, USA) according to the manufacturer’s protocol. Cells were transfected with FoxO1 and FoxO3a expression plasmids ^53^, *Ass1*-enhancer firefly luciferase reporter plasmids and Renilla luciferase reporter plasmid (pRL-SV40; Promega, Madison, WI, USA) after adjusting total amounts of transfected DNA with empty plasmid. After 48 hours cells were lysed with 100 μL of Reporter Lysis Buffer (Cat# E397A, Promega, Madison, WI, USA) and centrifuged. The supernatant was mixed with a luminometer with a firefly luciferase assay reagent (Cat# PGL5500, Pikkagene, Toyo Bnet bio, Tokyo, Japan) and the firefly luciferase activity was measured using a Wallac ARVO SX 1420 luminometer (PerkinElmer, Shelton, CT, USA). Renilla luciferase activity was measured with Renilla Luciferase Assay System (Cat# E2820, Promega, Madison, WI, USA) according to the manufacturer protocol. The Renilla luciferase activities were used to normalize transfection efficiencies.

### Electrophoretic mobility shift assay (EMSA)

EMSA were performed as described previously ^71^. In brief, the DNA probes were prepared by annealing two oligonucleotides, labeled with [α-^32^P] dCTP by filling in the 5’-overhangs with Klenow DNA polymerase, and purified on Sephadex G-50 columns. EMSA probes are listed in Table S3. A DNA fragment encoding the amino acid sequence (157 aa to 268 aa) containing the mouse FoxO1 DNA binding domain was inserted into the MCS of pGEX-4T1 to prepare a GST-fusion protein (GST-FoxO1-DBD). The labeled DNA probes were incubated with GST and GST-FoxO1-DBD in binding buffer (10 mM HEPES at pH 7.8, 50 mM KCl, 1 mM EDTA, 5 mM MgCl_2_, 10% glycerol, 5 mM dithiothreitol, and 0.4 μg/mL poly(dI-dC)), for 30 min on ice. The DNA-protein complex was analyzed on 4.6% polyacrylamide gels in TBE buffer.

GST and GST fusion proteins were expressed in *E. coli* (DH5α) using pGEX-4T (Amersham BioSciences, Buckinghamshire, UK) and purified using glutathione-Sepharose beads (Amersham BioSciences, Buckinghamshire, UK) by standard method as described previously ^36^.

### Preparation and transduction of recombinant adenoviruses

The DNA fragment of mouse *Ass1* enhancer region containing the FoxO binding site was amplified by PCR using PrimeSTAR GXL DNA Polymerase (Cat#R050A, TaKaRa Bio, Tokyo, Japan), using mouse genomic DNA as template and inserted into multiple cloning site on the pENTR4-Luc Gateway entry vector linked to firefly luciferase reporter with *Ass1* native promoter (*Ass1*-Enhancer-Luc) by infusion technique using In-Fusion HD Cloning Kit (Cat#071320, TaKaRa Bio, Tokyo, Japan) according to the manufacturer’s protocol. Mutated *Ass1*-Enhancer-Luc plasmid was generated using PrimeSTAR Mutagenesis Basal Kit (Cat# R046A, TaKaRa Bio, Tokyo, Japan). Primer sets are listed in Table S2. Adenoviral constructs were generated by homologous recombination between the entry vector and the pAd promoterless vector (Thermo Fisher Scientific, Waltham, MA, USA). Dominant-negative form FoxO1/3/4 (FoxODN), adenovirus encoding GFP (Ad-GFP), *FoxO*i and *LacZ*i, the two mouse *Ass1*-specific shRNA constructs (*Ass1*i-1, *Ass1*i-2) construction was described previously ^38,47^. Recombinant adenoviruses were propagated in HEK293 cells. Recombinant adenoviruses were then purified by CsCl gradient centrifugation, and titers were quantified as described previously ^36,38,50,72^. For animal experiments, adenoviruses were injected intravenously into ICR male mice from subclavian vein at the following doses: for *LacZ*i/*FoxO*i, 10x10^8 P.F.U.; for GFP/FoxO-DN, 5x10^8 P.F.U.; *Ass1*-enhancer-Luc, 2x 10^8 P.F.U.; *LacZ*i/*Ass1*i 10x10^8 P.F.U. (1 P.F.U. is equal to 1000 optical particles of adenovirus).

### In vivo imaging of Luciferase Activity

*In vivo* imaging was performed as described previously ^36,50–52^. After 4 days of the adenovirus transduction, animals were fasted for 24-h from the early dark phase. At each condition, D-Luciferin potassium salt (Cat# 126-05116, Wako Chemicals, Tokyo, Japan) dissolved in PBS at a concentration of 7.5 mg/mL was intraperitoneally injected at a dose of 10 mL/kg into mice and the luminescence in the liver was captured using an IVIS^TM^ Imaging System (PerkinElmer, Waltham, MA, USA). Relative photon emissions over the liver region was quantified using LIVING IMAGE^TM^ software (PerkinElmer). Hepatic transduction efficiency was determined based on the quantification of adenoviral DNA in the liver using a previously described Q-PCR method ^72^, and the result of quantification was used to normalize the in vivo imaging of luciferase activity. Two paired data from the same animal on different nutritional conditions (i.e., before and after fasting) were obtained and the ratio between the two quantities was used to cancel the variations in hepatic transduction efficiencies.

### Chromatin immunoprecipitation (ChIP) assay

Chromatin immunoprecipitation (ChIP) assays using mouse liver were performed as described previously ^38,47^. Briefly, 100 mg of liver tissue samples from the fasted and ad libitum-fed mice were minced in 1 mL of PBS and cross-linked in 1.5% formaldehyde for 15 min at room temperature. Fixed samples were homogenized and then subjected to sonication with Bioruptor2 (Sonicbio, Kanagawa, Japan) for DNA fragmentation. After centrifugation, supernatant was diluted to 6 mL with dilution buffer (50 mM Tris-HCl at pH 8.0, 167 mM NaCl, 1 mM EDTA, 1.1% Triton X-100, and 0.1% sodium deoxycholate) and then 1 mL of total volume was used for immunoprecipitation with 1 μg of anti-FoxO1 (Cat#2880, Cell Signaling Technology), anti-FoxO3a (Cat#12829, Cell Signaling Technology), or control IgG (Cat# CR1, Sino Biological Inc.) bound to 30 μL of Dynabeads magnetic beads (Thermo Fisher Scientific, Waltham, MA, USA) and rotated overnight at 4 °C. Fifty microliter of total volume was used for input sample. The complexes were washed with low-salt wash buffer (50 mM Tris-HCl at pH 8.0, 150 mM NaCl, 1 mM EDTA, 1% Triton X-100, 0.1% SDS, and 0.1% sodium deoxycholate), high-salt wash buffer (50 mM Tris-HCl at pH 8.0, 500 mM NaCl, 1 mM EDTA, 1% Triton X-100, 0.1% SDS, and 0.1% sodium deoxycholate), LiCl wash buffer (10 mM Tris-HCl at pH 8.0, 0.25 M LiCl, 1 mM EDTA, 0.5% NP40, and 0.5% sodium deoxycholate), and TE buffer. DNA-protein complex was eluted by incubation with elution buffer (1% SDS, 0.1 M NaHCO_3_) for 15 min at room temperature and then incubated with 200 mM NaCl overnight at 65 °C for reverse crosslinking. DNA-protein complex was treated with 200 μg/mL proteinase K, and chromatin DNA was purified with phenol-chloroform, eluted in TE buffer, and subjected to Q-PCR analysis. From reverse crosslinking step, the input sample was also subjected to the same procedure. Q-PCR was performed using the same method as Q-RT PCR and quantified by standard curve method with input DNA samples. The primer sets are listed in Table S4.

### Total liver lysate

For total liver lysate, 20 mg of liver tissues collected from 3-4 mice were pooled and homogenized in 1 mL lysis buffer (50 mM Tris-HCl at pH 7.5, 137 mM NaCl, 1 mM EDTA, 1% Triton-X, protease inhibitors). After centrifugation at 15,000 rpm for 10 min at 4 °C, the supernatant was used as a total liver lysate.

### Liver nuclear extraction

Liver nuclear extraction was performed as previously described ^73^. In brief, 1.5 g of liver tissues collected from 3-4 mice were pooled and homogenized in 15 mL of buffer A (10 mM HEPES at pH 7.6, 25 mM KCl, 1 mM EDTA, 2 M sucrose, 10% glycerol, 0.15 mM spermine, 2 mM spermidine, protease inhibitors). The sample was filtered with sterile gauze (Kawamoto Corporation) and layered on 15 mL of buffer A in a polypropylene centrifuge tube (Beckman coulter, Brea, CA, USA). The tube was centrifuged at 24,000 rpm for 90 min at 4 °C. The pellet was suspended in 800 μL of buffer B (10 mM HEPES at pH 7.6, 100 mM KCl, 2 mM MgCl_2_, 1 mM EDTA, 1 mM DTT, 10% glycerol, protease inhibitors) and centrifuged at 89,000 rpm for 20 min at 4 °C. The supernatant was used as a nuclear extract.

### Western Blotting

Western blotting was performed as previously described ^74,75^. In brief, after SDS-PAGE, proteins were transferred onto nitrocellulose membrane (Hybond ECL, Amersham BioSciences, Buckinghamshire, UK). Ass1, GAPDH, FoxO1, FoxO3a, and Lamin were detected using a 1:1000 dilution mouse anti-Ass1 (Cat#sc-365475, Santa Cruz Biotechnology, Dallas, TX, USA), and mouse anti-GAPDH (Cat#sc-32233, Santa Cruz Biotechnology, Dallas, TX, USA), rabbit anti-FoxO1 (Cat#2880, Cell Signaling Technology, Danvers, MA, USA) and rabbit anti-FoxO3a (Cat#2497, Cell Signaling Technology, Danvers, MA, USA), and mouse anti-LaminA/C (Cat#sc-376248, Santa Cruz Biotechnology, Dallas, TX, USA), in TBS buffer (20 mM Tris-HCl at pH 7.6 and 140 mM NaCl) containing 0.2% Tween 20 and 5% skimmed milk. Bound antibodies were detected with a horseradish peroxidase-coupled anti-mouse IgG secondary antibody (Cat#7076, Cell Signaling Technology, Danvers, MA, USA) or anti-rabbit IgG secondary antibody (Cat#7074, Cell Signaling Technology, Danvers, MA, USA) and visualized using anti-ECL chemiluminescent substrates (Cat# NEL104001EA, Revvity Health Sciences Inc.).

### Statistical analysis

Data are expressed as means ± S.E.M. Differences between two groups were assessed using the unpaired two-tailed Student’s *t*-test and the paired two-tailed Student’s *t*-test for IVIS data analysis. The differences were considered to be significant if *P* < 0.05. (\**P* < 0.05 and \*\**P* < 0.01).

## Supplementary Information

**Figure S1.**
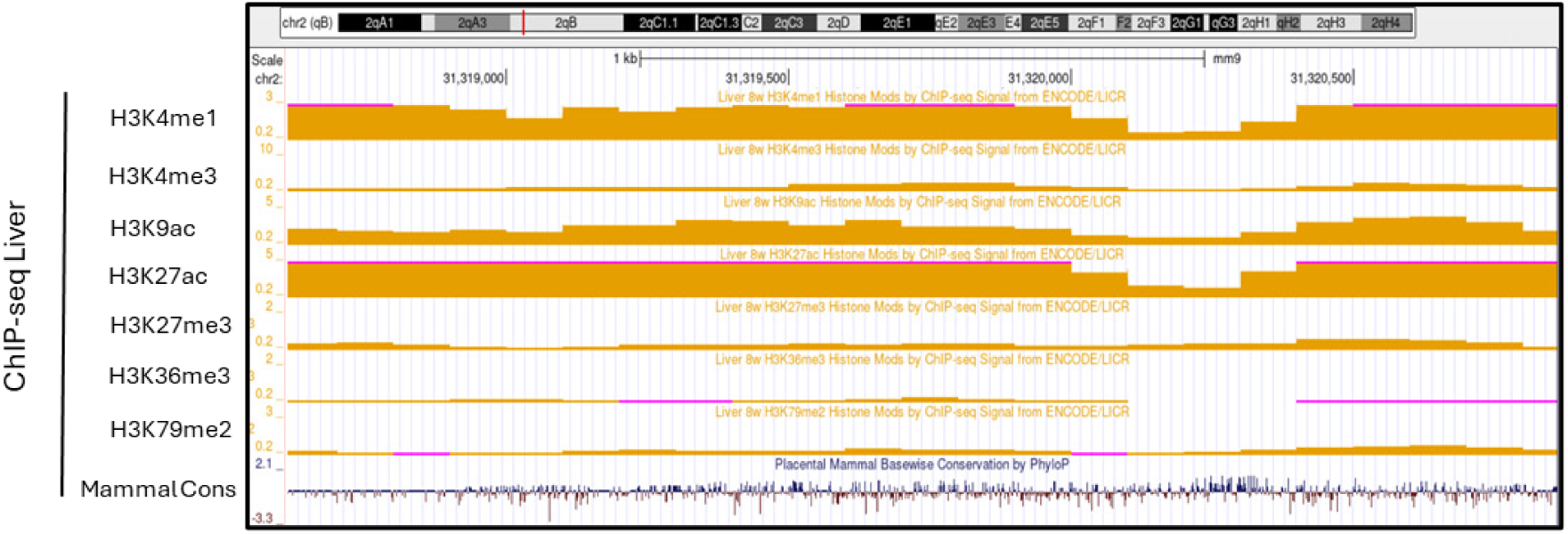
Genomic information on *Ass1* enhancer. Modified Histones binding regions on *Ass1 enhancer* from ENCODE ChIP-seq database (Liver 8w by ChIP-seq Signal from ENCODE/LICR) on Genome browser. These peaks mean promoter/enhancer regions. Mammalian conservation regions are shown.

**Figure S2.**
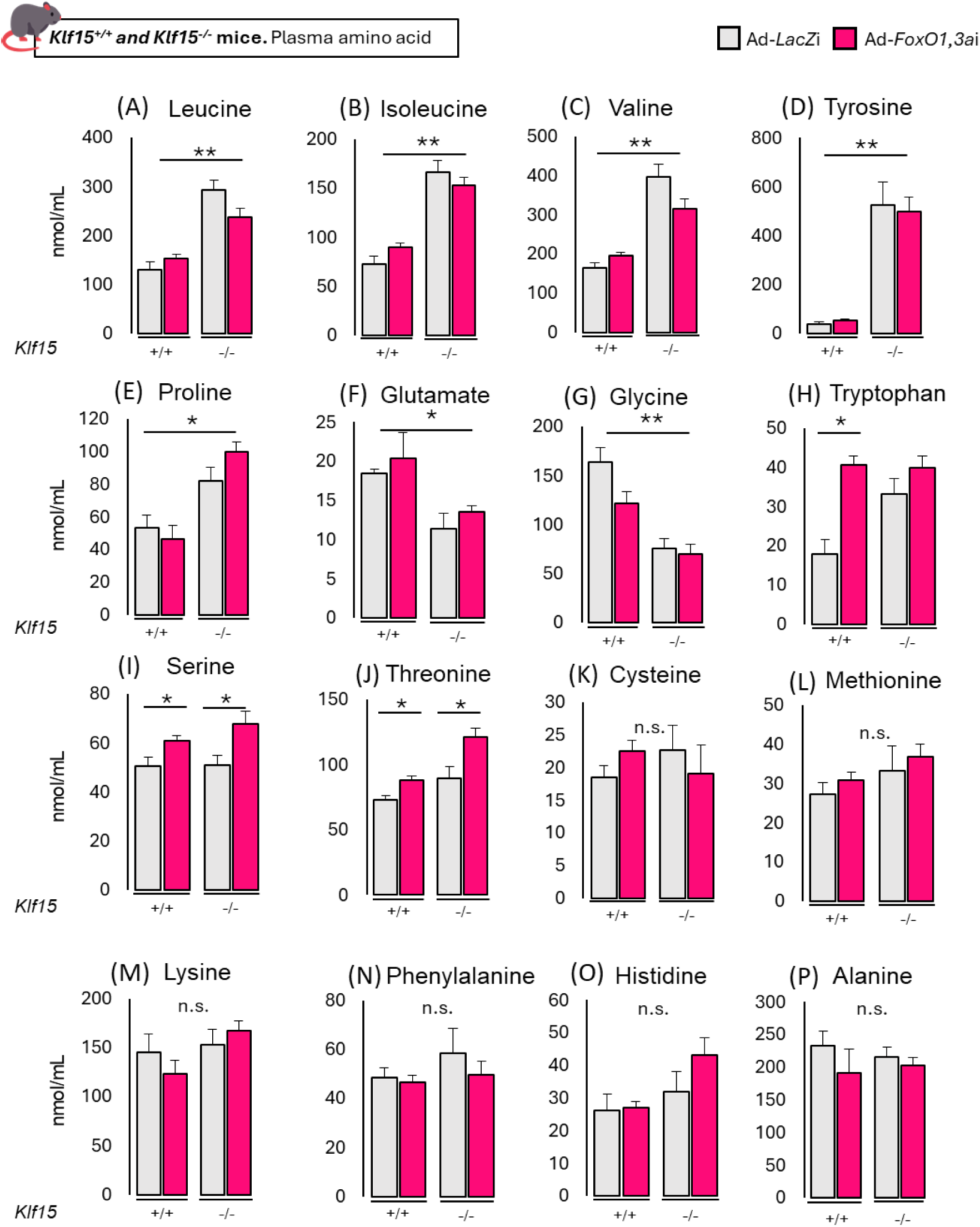
Plasma amino acids profile in *Klf15* wild-type and knockout mice. Amino acid levels in the plasma of *Klf15* wild-type and knockout C57BL/6J mice with hepatic *FoxO1*/*FoxO3a* knockdown (n = 4 per group). *FoxO1* and *FoxO3a* knockdown were performed using adenovirus mediated RNAi (Ad-*FoxO1,3a*i). All groups of mice were sacrificed in the light cycle and blood samples were collected in the fasted state following 24-h fasting. All values are presented as the mean with error bars representing the SEM. Datasets were assessed by Student’s *t*-test for unpaired samples. The differences were considered to be significant if *P* < 0.05 (\**P* < 0.05 and \*\**P* <0.01).

**Figure S3.**
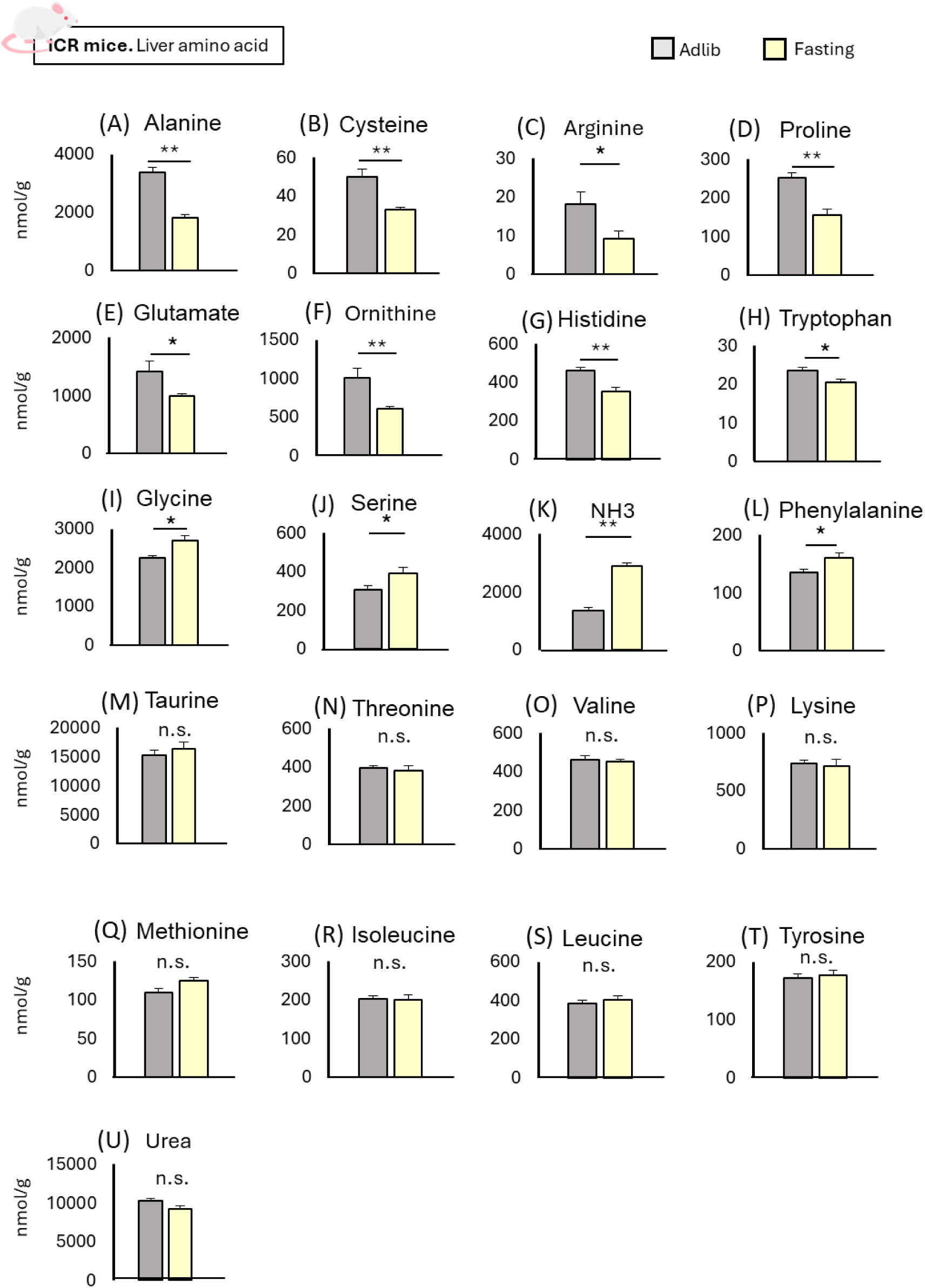
Hepatic amino acid profile in ICR mice during fasting. Amino acid levels in the liver of ICR mice in ad libitum and fasted state (n = 6 per group). All groups of mice were sacrificed in the light cycle and liver samples were collected in the fasted state following 24-h fasting or ad libitum as indicated. All values are presented as the mean with error bars representing the SEM. Datasets were assessed by Student’s *t*- test for unpaired samples. The differences were considered to be significant if *P* < 0.05 (\**P* < 0.05 and \*\**P* <0.01).

**Figure S4.**
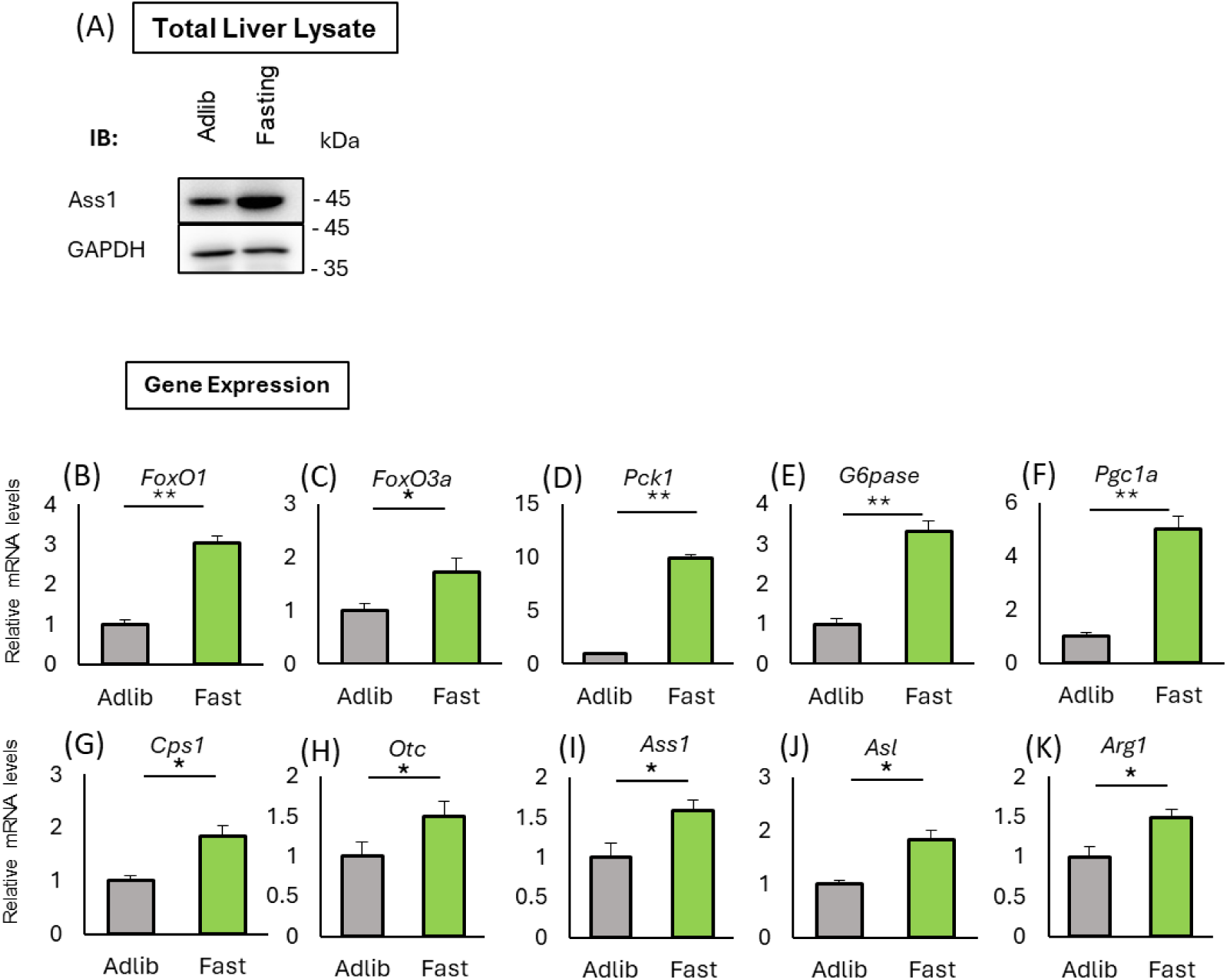
Effect of fasting on FoxO transcription factors, gluconeogenesis and urea cycle genes. (A) Immunoblot analysis of Ass1 protein using liver total lysate from mice in the fasted and ad libitum state (Samples were pooled from 3 mice). GAPDH was detected as an internal loading control. (B-K) Q-RT PCR analysis of liver RNA samples. Relative gene expression was analyzed in ICR mice in ad libitum and fasted state (n = 6 per group). As the correction of the gene expression level for each sample *Cyclophilin A* was used. All groups of mice were sacrificed in the light cycle and liver samples were collected in and the fasted state following 24-h fasting or ad libitum as indicated. All values are presented as the mean with error bars representing the SEM. Datasets were assessed by Student’s *t*- test for unpaired samples. The differences were considered to be significant if *P* < 0.05 (\**P* < 0.05 and \*\**P* <0.01).

**Figure S5.**
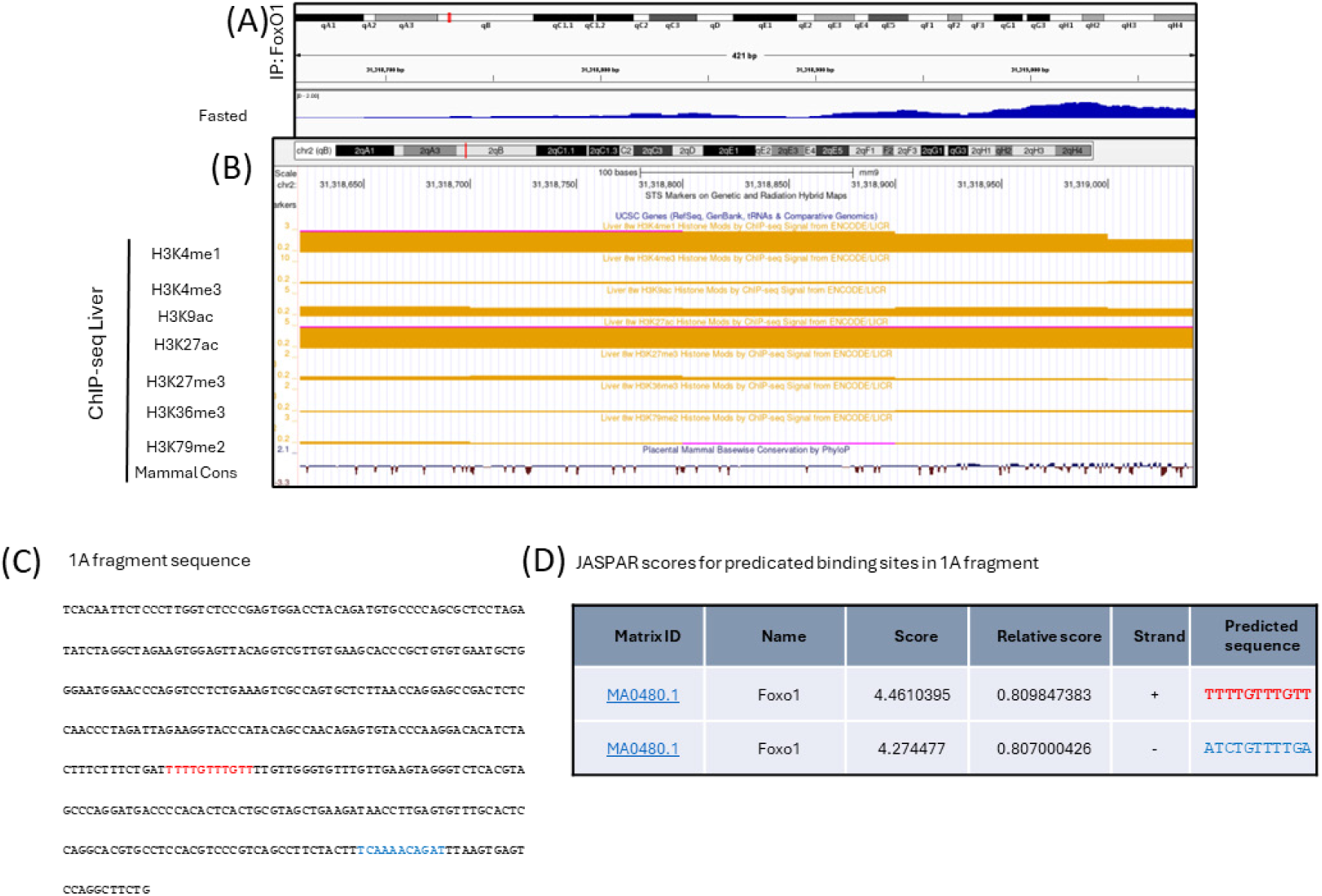
Genomic information on FoxO binding site in the *Ass1* enhancer, related to. Figure 4. (A) the results of ChIP-seq with anti-FoxO1 antibody in livers of unfed mice obtained from ChIP-Atlas (ID: GSM3381273) of the smallest fragment 1A. (B) Modified Histones binding regions on *Ass1* enhancer from ENCODE ChIP-seq database (Liver 8w by ChIP-seq Signal from ENCODE/LICR) on Genome browser. These peaks means promoter/enhancer regions. Mammalian conservation regions are shown below ChIP-seq peaks data. (C) Mouse genomic sequence of the smallest fragment 1A. (D) Estimation of FoxOs binding sites on the smallest fragment 1A by JASPAR2020 (http://jaspar.genereg.net/). In red is *Ass1* FoxO BS1, in blue is *Ass1* FoxO BS2 were used as probs for EMSA in Figure 4D and 4E.

**Table S1.**
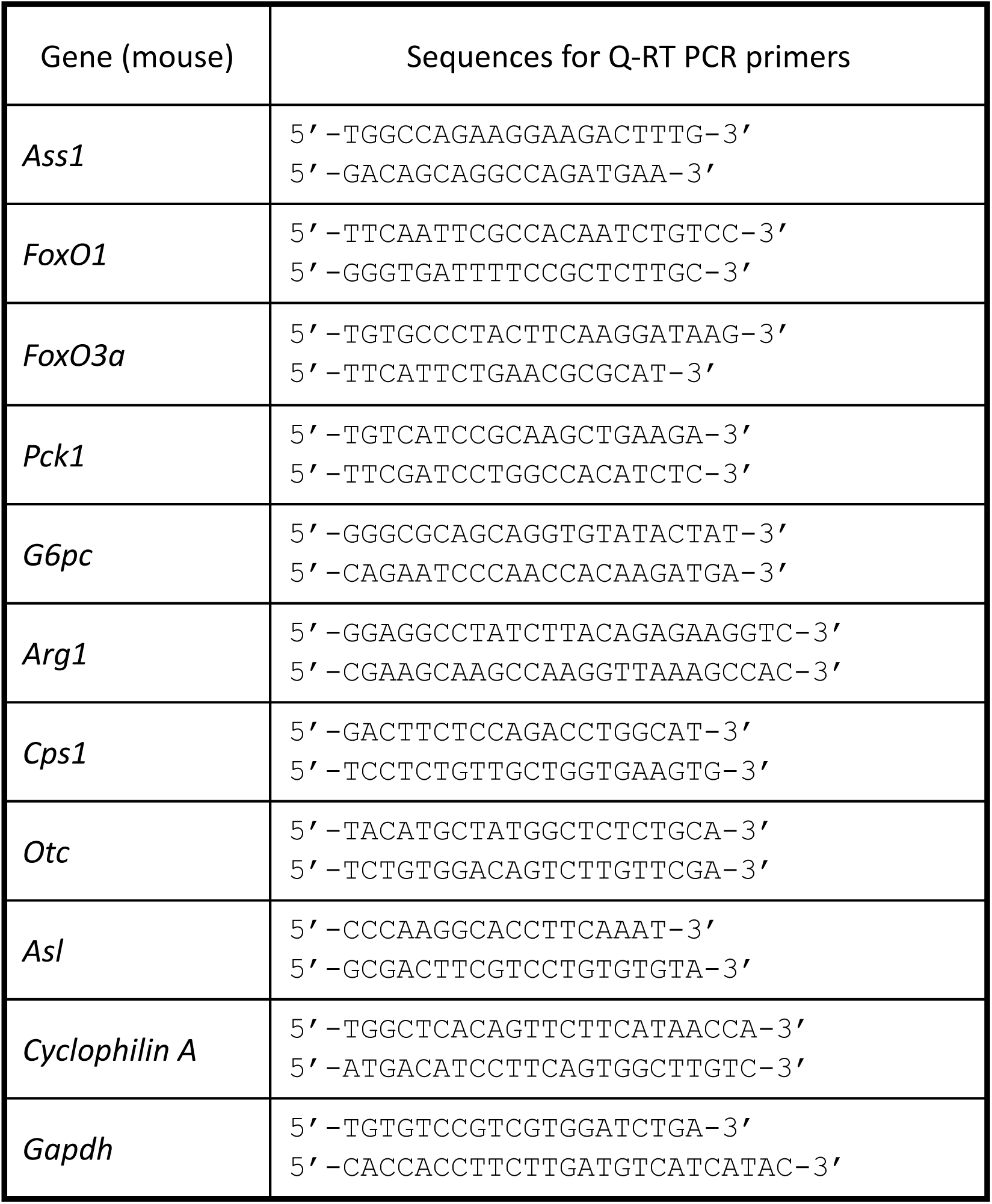
List of primer sets used for Q-RT PCR.

**Table S2.**
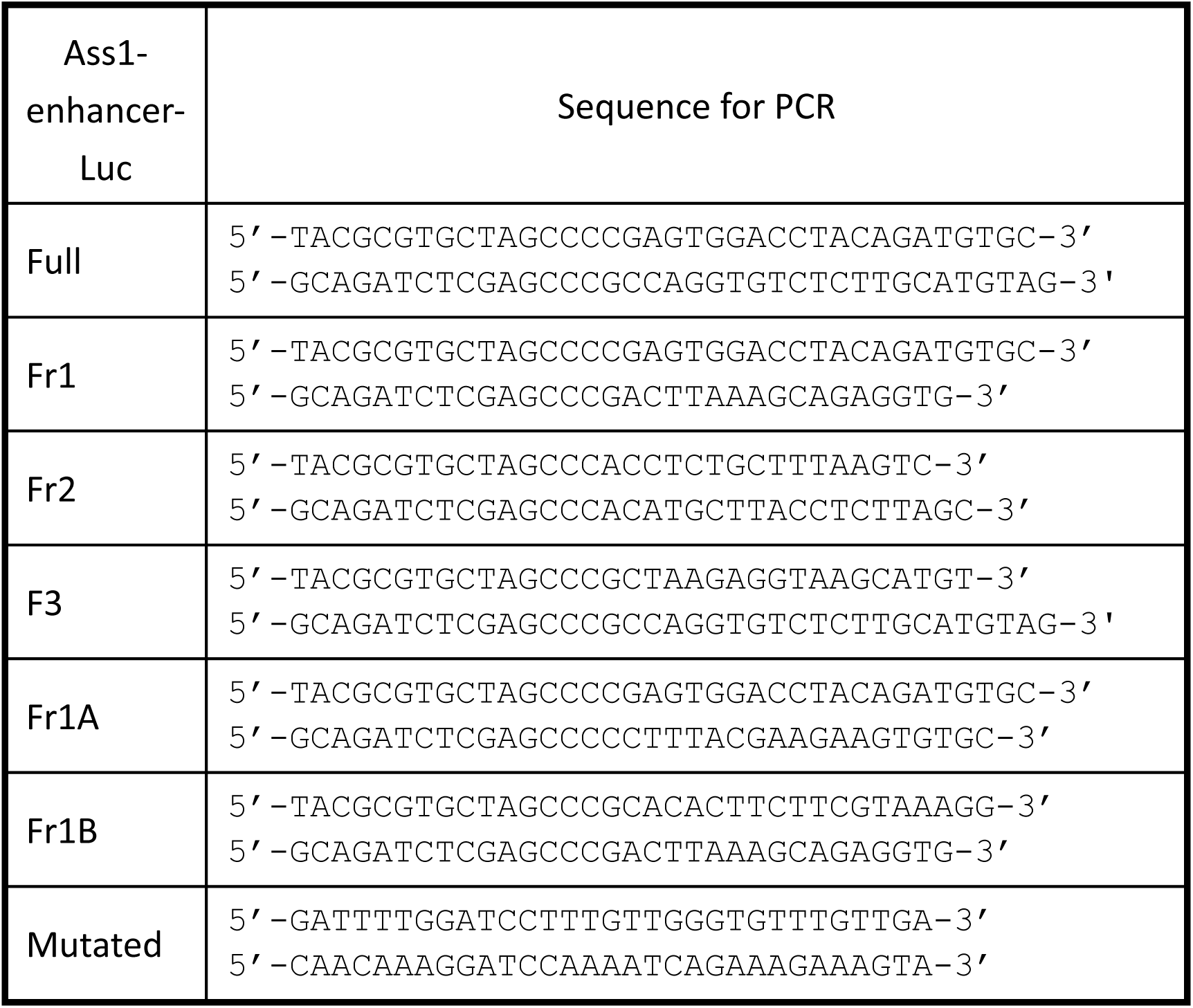
List of primer sets used for PCR.

**Table S3.**
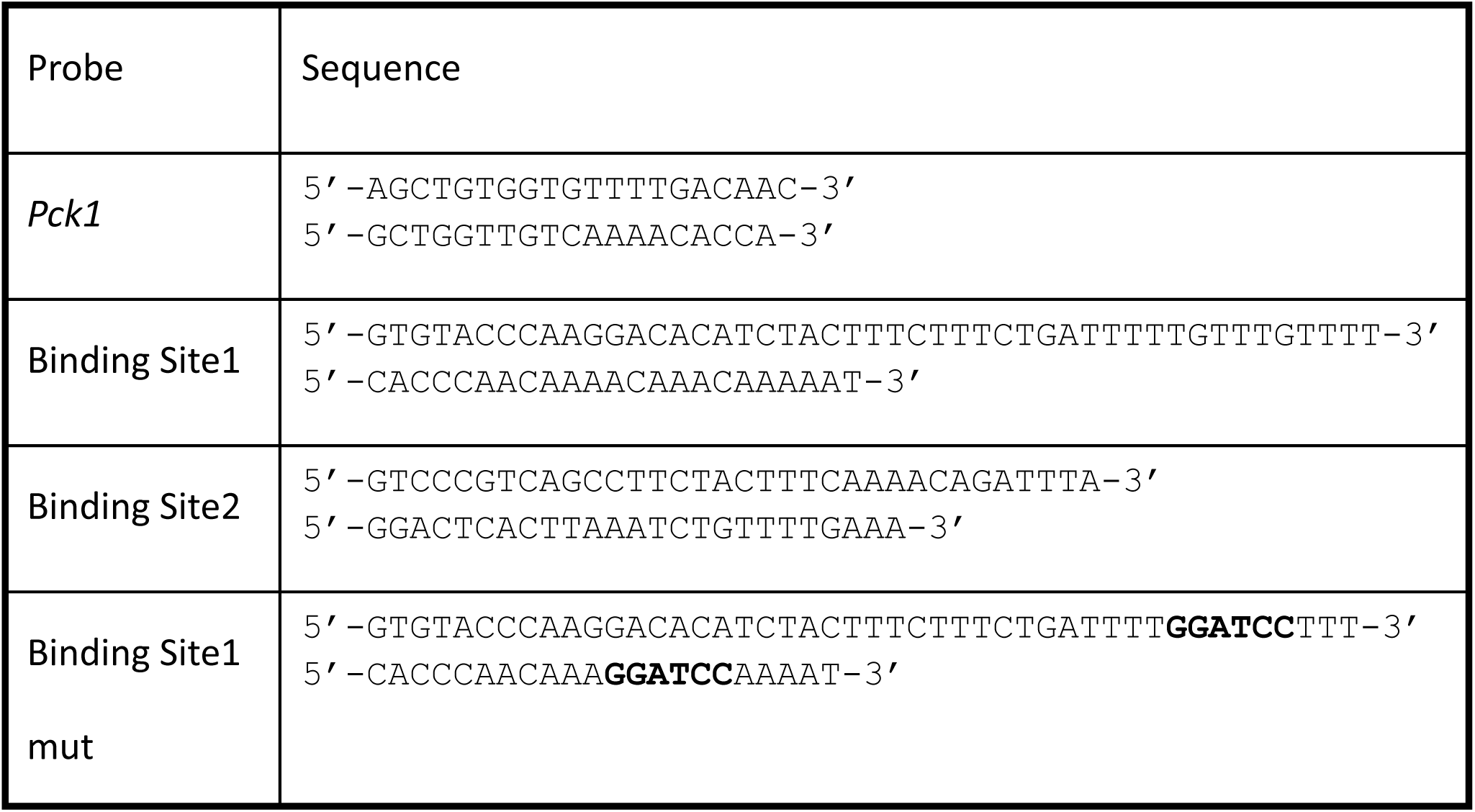
List of EMSA probes.

**Table S4.**
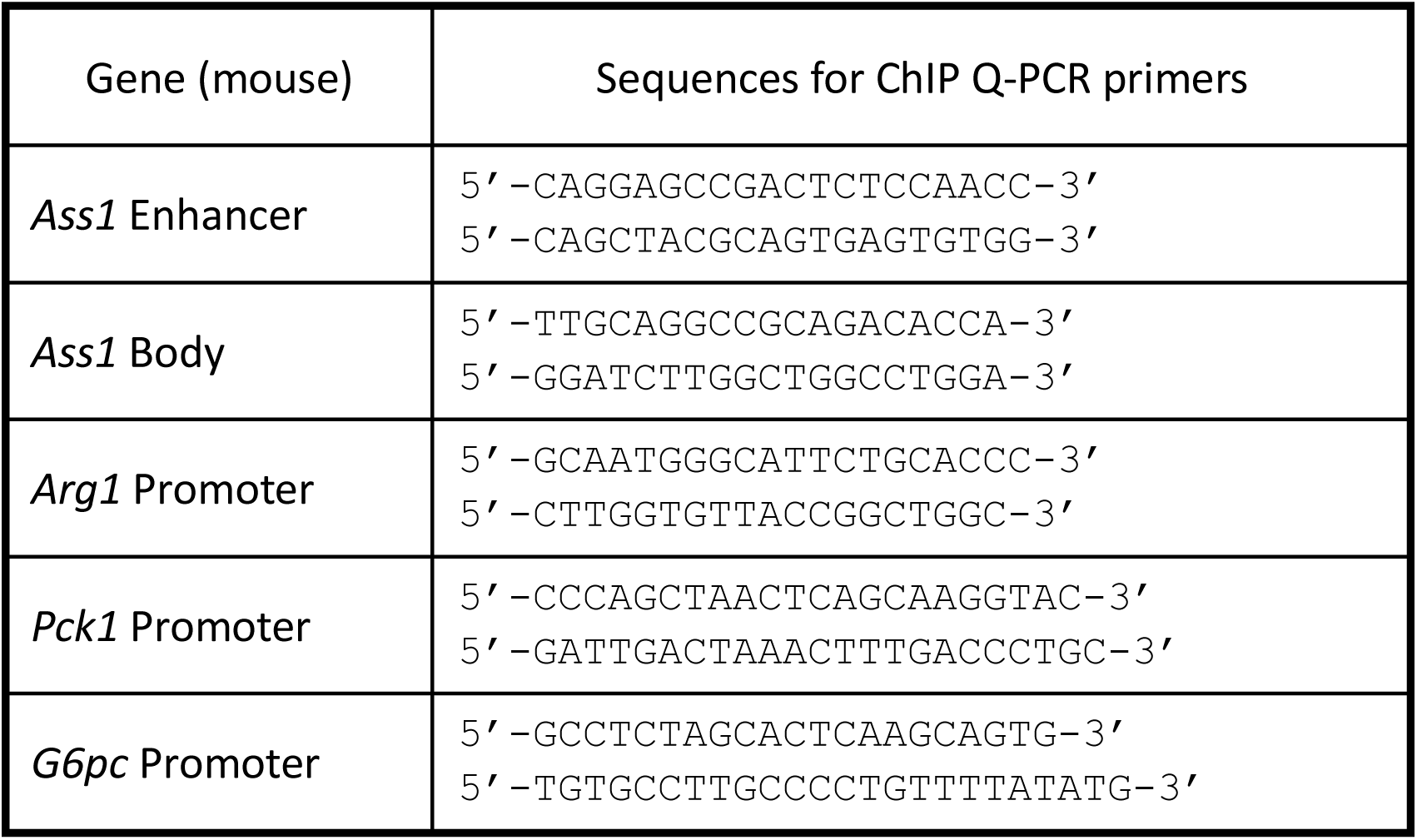
List of primer sets used for ChIP assay.

